# The plant nucleoplasmin AtFKBP43 needs its extended arms for histone interaction

**DOI:** 10.1101/2022.04.25.489347

**Authors:** Ajit Kumar Singh, Ketul Saharan, Somanath Baral, Dileep Vasudevan

**Affiliations:** Institute of Life Sciences, Bhubaneswar, India 751023; Regional Centre for Biotechnology, Faridabad, India 121001; School of Biotechnology, KIIT University, Bhubaneswar, India 751024

**Keywords:** Nucleoplasmin, FKBP, Histone chaperone, FKBP nucleoplasmin, Crystal structure, SAXS

## Abstract

The nucleoplasmin family of histone chaperones is a key player in governing the dynamic architecture of chromatin, thereby regulating various DNA-templated processes. The crystal structure of the N-terminal domain of *Arabidopsis thaliana* FKBP43 (AtFKBP43), an FK506-binding immunophilin protein, revealed a characteristic nucleoplasmin fold, thus confirming it to be a member of the FKBP nucleoplasmin class. Small-Angle X-ray Scattering (SAXS) analyses confirmed its pentameric nature in solution, and additional studies confirmed the nucleoplasmin fold to be highly stable. The AtFKBP43 nucleoplasmin core domain could not interact with histones and required the acidic arms, C-terminal to the core, for histone association. Furthermore, SAXS generated low-resolution envelope structure, ITC, and AUC results revealed that an AtFKBP43 pentamer with C-terminal extensions interacts with H2A/H2B dimer and H3/H4 tetramer in an equimolar ratio. Put together, AtFKBP43 belongs to a hitherto unreported subclass of FKBP nucleoplasmins that requires the C-terminal acidic stretches emanating from the core domain for histone interaction.

## INTRODUCTION

The core of a nucleosome, the basic repeating unit of chromatin in eukaryotes (1, 2), consists of a negatively charged ∼147 bp DNA wrapped around a positively charged histone octamer (3, 4). The nucleosome must undergo assembly and disassembly to allow access to the DNA-templated functions (5-7). These dynamic nucleosome events are regulated by factors such as chromatin remodelers and histone chaperones (8, 9). The different families of histone chaperones are important players, carrying out diverse functions such as preventing the aggregation and non-specific interaction of histones with DNA and other cellular components (10, 11), aiding in folding, storing, and transporting histones from the cytoplasm to the nucleus (12, 13). Histone chaperones also facilitate the transfer of histones onto the DNA (deposition) and off from the DNA (eviction), thereby regulating various dynamic chromatin processes (14)

The nucleoplasmin (NPM) family of histone chaperones is found throughout the animal kingdom in somatic and germinal cells (15-17). They generally exist as a pentamer in solution, forming a decamer in a few instances (18). Common to the members of the nucleoplasmin family is a well-conserved protease-resistant, thermally stable N-terminal oligomerization domain or the core domain (19), which also includes, in most cases, a short A1 acidic tract. The intrinsically disordered protease-sensitive C-terminal domain/tail of variable length, containing two acidic tracts, namely A2 and A3, emanates from the core domain protomers, often plays a role in histone binding and charge shielding, though the core alone is enough for histone interaction (20, 21)

Nucleoplasmins have been categorized into four main groups: Nucleophosmin (NPM1), Nucleoplasmin (NPM2), Nucleoplasmin-like (NPM3), and invertebrate NPM based on protein sequences.^15^ The crystal structure of the core domains of several nucleoplasmins has been reported. The structures reveal that each monomer possesses an eight-stranded β-sandwich fold and that five such monomers associate to form a cyclic pentamer (20). According to a more recent classification, nucleoplasmins are divided into (i) classical nucleoplasmins, (ii) FKBP nucleoplasmins (those possessing a C-terminal FK506-binding domain, in addition to the N-terminal nucleoplasmin core; present in yeast, insects, and plants), and (iii) HDT-nucleoplasmins (those belonging to deacetylase class and present only in plants) (19, 22). Phylogenetic analyses have suggested the presence of FKBP nucleoplasmins in various organisms such as yeasts, arthropods, and plants, but it is absent among vertebrates (19). FKBPs are members of the larger family of immunophilins and form the binding targets for the immunosuppressant drugs FK506 and rapamycin (23). In addition, they catalyze the cis-trans interconversion of peptidyl-prolyl bonds, which helps properly fold a nascent protein (24). Immunophilins being a major group of protein folding catalysts, few of the nuclear-localized multi-domain FKBPs (a member of immunophilin family proteins), such as *Saccharomyces cerevisiae* Fpr3 and Fpr4 (ScFPR3 and ScFPR4), *Schizosaccharomyces pombe* FKBP39 (SpFKBP39), and *Drosophila melanogaster* FKBP39 (DmFKBP39) have recently been reported to play a role in chromatin remodeling (25-27). The multi-domain FKBPs have additional copies of FKBDs or other functional domains (28), such as the FKBP nucleoplasmins that possess an N-terminal nucleoplasmin domain (19). Among the several FKBP nucleoplasmins, structural information is available for the nucleoplasmin domain of DmFKBP39 (19) and both the domains of AtFKBP53 (29). AtFKBP53 has also been identified to interact with histones (30, 31) and has been confirmed to be a functional histone chaperone, aiding nucleosome assembly (29). AtFKBP43 has been proposed to be a close homolog of AtFKPB53 with possibly overlapping functional attributes (30). However, the protein has so far not been characterized. Herein, we report the structural and functional features of AtFKBP43.

## RESULTS

### AtFKBP43 appears to be a nucleoplasmin according to sequence, phylogenetic and secondary structure analyses

The domain organization of AtFKBP43 revealed an acidic domain at its N-terminus (1-96) and a basic peptidyl-prolyl cis-trans isomerase (PPIase) or FK506-binding domain (FKBD) at its C-terminus (350-499) predicted to take up ordered conformations, whereas the middle region (110-383) appeared highly disordered **(Fig. 1A and Fig. S1A)**. Nucleoplasmins possess a C-terminal stretch of variable length with multiple acidic tracts (32). The AtFKBP43 sequence revealed four acidic stretches, namely A1 (residues 73-79), A2 (residues 107-136), A3 (residues 138-164) and A4 (residues 186-200) **(Fig. 1A)**. Similar to other nucleoplasmins, A2 is the longest acidic tract in AtFKBP43.

**Figure 1.**
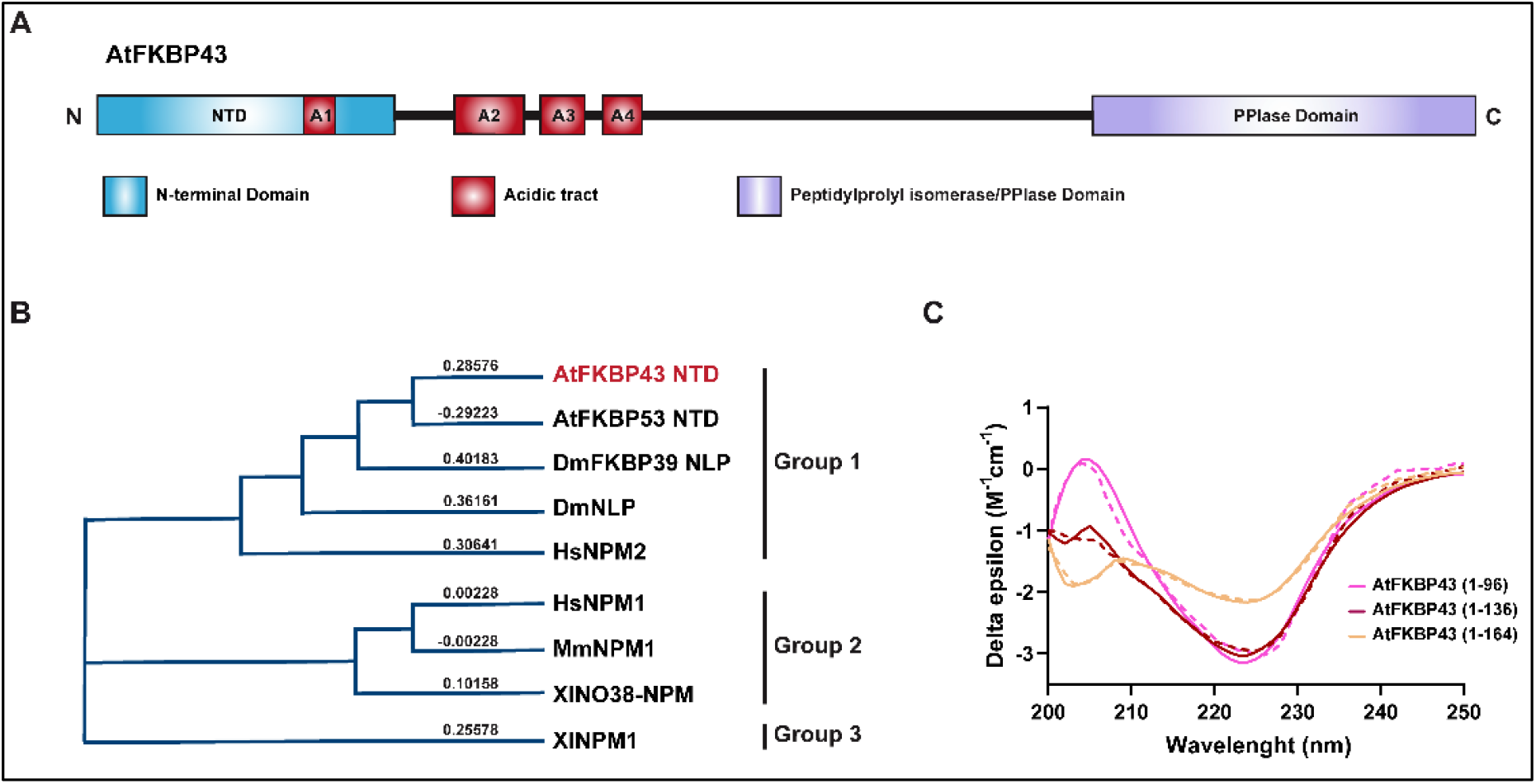
Domain organization, phylogenetic, and CD analysis of AtFKBP43. **(A)** A representation of the domain organization in AtFKBP43. NTD is shown in cyan, acidic tracts are shown in red, and the PPIase domain in violet. The acidic tracts are labelled A1 to A4. **(B)** Phylogenetic tree constructed based on T-Coffee sequence alignment program, using the Neighbour-Joining method for AtFKBP43 NTD, AtFKBP53 NTD, DmFKBP39 NLP, XlNPM, HsNPM1, MmNPM1, HsNPM2, XlNO38 NPM and DmNLP. The values shown against the proteins represent the evolutionary distance between the sequences. **(C)** The CD spectrum of AtFKBP43 NTDs. The CD curves are shown for AtFKBP43 (1-96) in pink, AtFKBP43 (1-136) in brown, and AtFKBP43 NTD (1-164) in sand yellow colour. The data deconvoluted by the online program BeStSel showed that the protein has a predominantly β-sheet conformation.

Multiple sequence alignment for AtFKBP43 NTD with other structurally characterized nucleoplasmins followed by phylogenetic analysis showed AtFKBP43 NTD is the closest to AtFKBP53 NTD and DmFKBP39-NLP with a sequence identity of 47.47% and 25.53%, respectively **(Fig. 1B and Fig. S1B)** and might take up a nucleoplasmin-like fold.

Further, CD spectroscopy displayed a high percentage of β-strands for AtFKKBP43 (1-96), (1-136) and (1-164) **(Fig. 1C and Table S1)**, similar to nucleoplasmin family proteins (33). These results together suggested AtFKBP43 to be a member of FKBP nucleoplasmin.

### AtFKBP43 NTD exists as a pentamer in solution

Earlier reports suggest that nucleoplasmins contain a well-folded N-terminal core domain that takes up a pentameric or a decameric conformation, followed by a C-terminal flexible region with distinct acidic stretches (20, 34). Analytical size-exclusion chromatography (analytical SEC) for AtFKBP43 NTD stretches 1-96, 1-136, and 1-164 gave homogenously distributed single peaks in their elution profiles **(Fig. 2A)**. However, AtFKBP43 NTD (1-200) appeared to form a soluble aggregate **(Fig. S2A)**. Further, sedimentation velocity analytical ultracentrifugation (SV-AUC) experiments for AtFKBP43 (1-96), (1-136), and (1-164) gave sedimentation coefficient (S) values of 3.90 S, 4.38 S, and 4.37 S that corresponds to a molecular mass of 57.08 kDa, 82.46 kDa and 97.62 kDa, respectively indicating a pentameric conformation for all **(Fig. 2B: Fig. S2, B, C, and D)**. Frictional ratio, Stokes radius, and Smax/S values **(Table 1)** suggested that AtFKBP43 (1-96) is relatively globular in shape, whereas AtFKBP43 (1-136) and (1-164) appeared elongated.

**Table 1:**
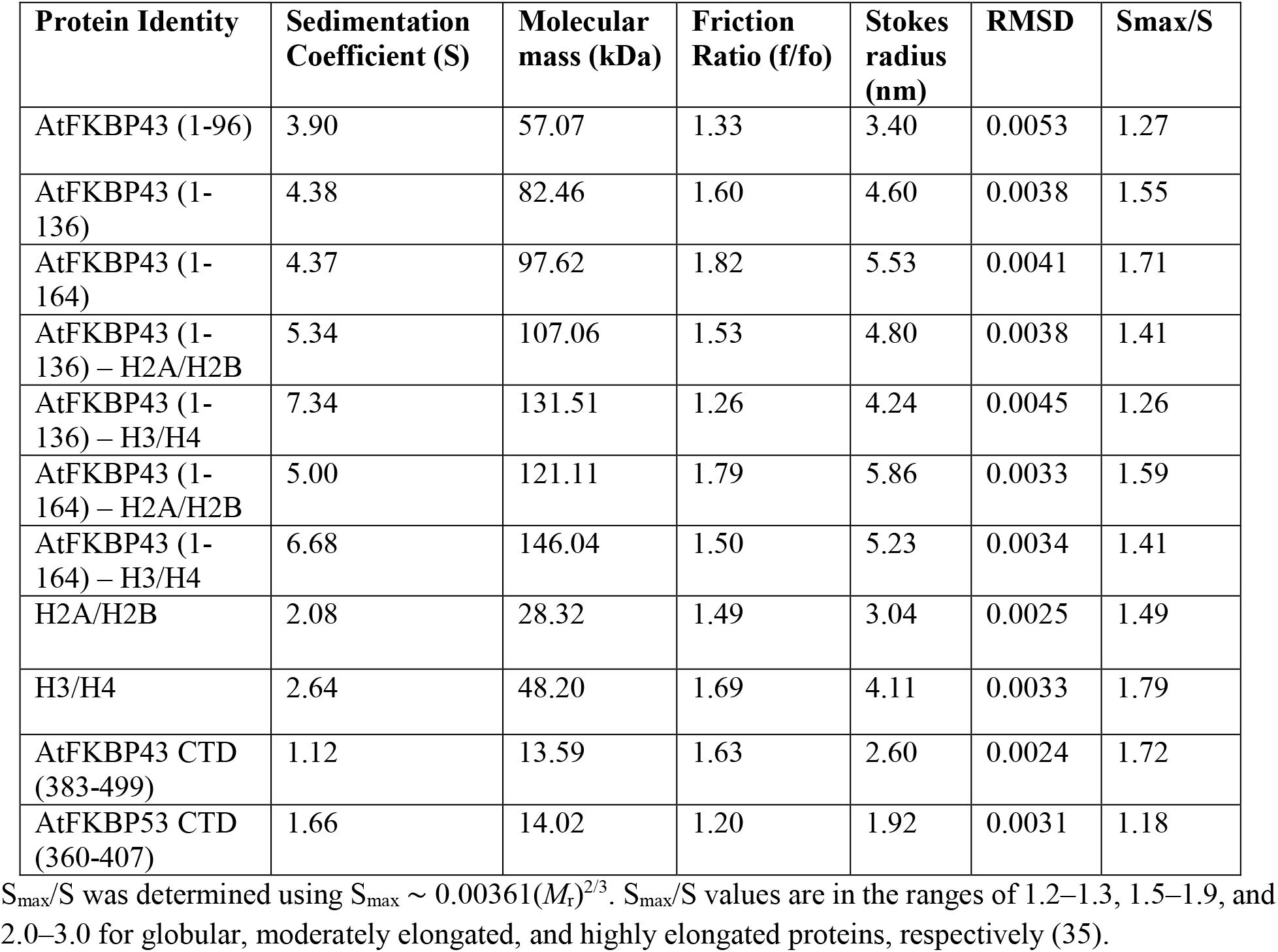
Sedimentation Velocity - Analytical Ultracentrifugation (SV-AUC) data for AtFKBP43 CTD, AtFKBP43 NTDs, histone oligomers, and their complexes.

**Figure 2.**
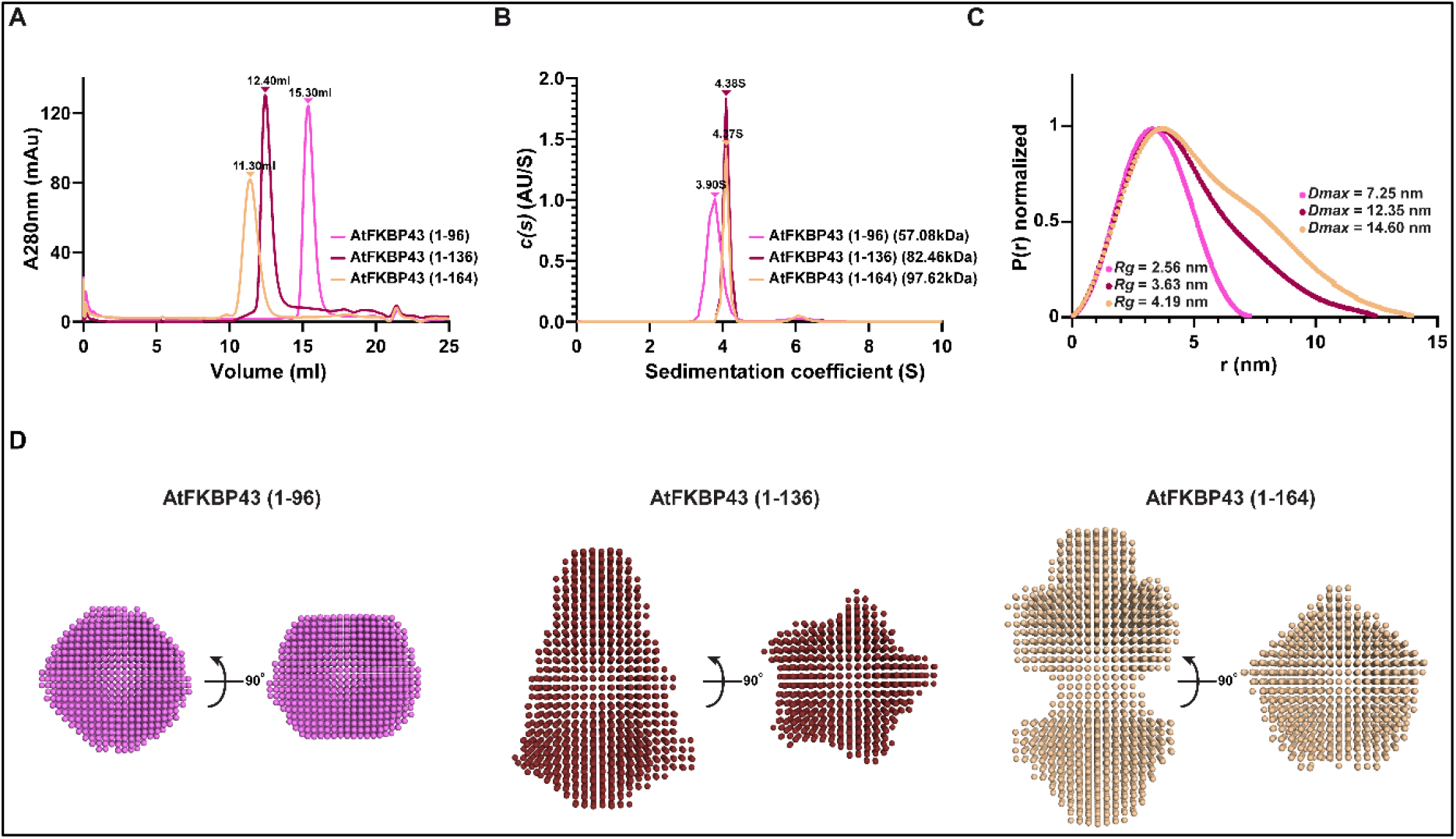
Oligomeric state analysis of AtFKBP43 NTDs. **(A)** Analytical SEC profiles of AtFKBP43 (1-96), (1-136), and (1-164). **(B)** SV-AUC statistical analysis of AtFKBP43 (1-96), (1-136), and (1-164). The figure shows the AUC distance distribution c(S) vs. sedimentation coefficient (S) plot obtained from SEDFIT software. **(C)** The Pairwise distribution function, P(r) of AtFKBP43 (1-96), (1-136) and (1-164). The real space Rg and Dmax values for AtFKBP43 (1-96), (1-136), and (1-164) are also shown. **(D)** The envelope structures for AtFKBP43 (1-96), (1-136), and (1-164), calculated from the SAXS data in P5 symmetry.

Further, the low-resolution structure of AtFKBP43 NTD stretches 1-96, 1-136, and 1-164 were determined using small-angle X-ray scattering (SAXS) experiments **(Fig. 2D)**. Details about data collection and other structural parameters have been reported in **Table 2**. The SAXS Guinier plot demonstrated AtFKBP43 NTDs as stable and aggregation-free **(Fig. S2E)**. The Kratky plot and Pairwise distribution function **(Fig. 2C and Fig. S2F)** confirmed a folded and globular nature for AtFKBP43 (1-96), whereas it suggested a partially folded and elongated conformation for AtFKBP43 (1-136) and (1-164). The real space Rg and Dmax calculated for the experimental data are provided in **Table 2**.

**Table 2:**
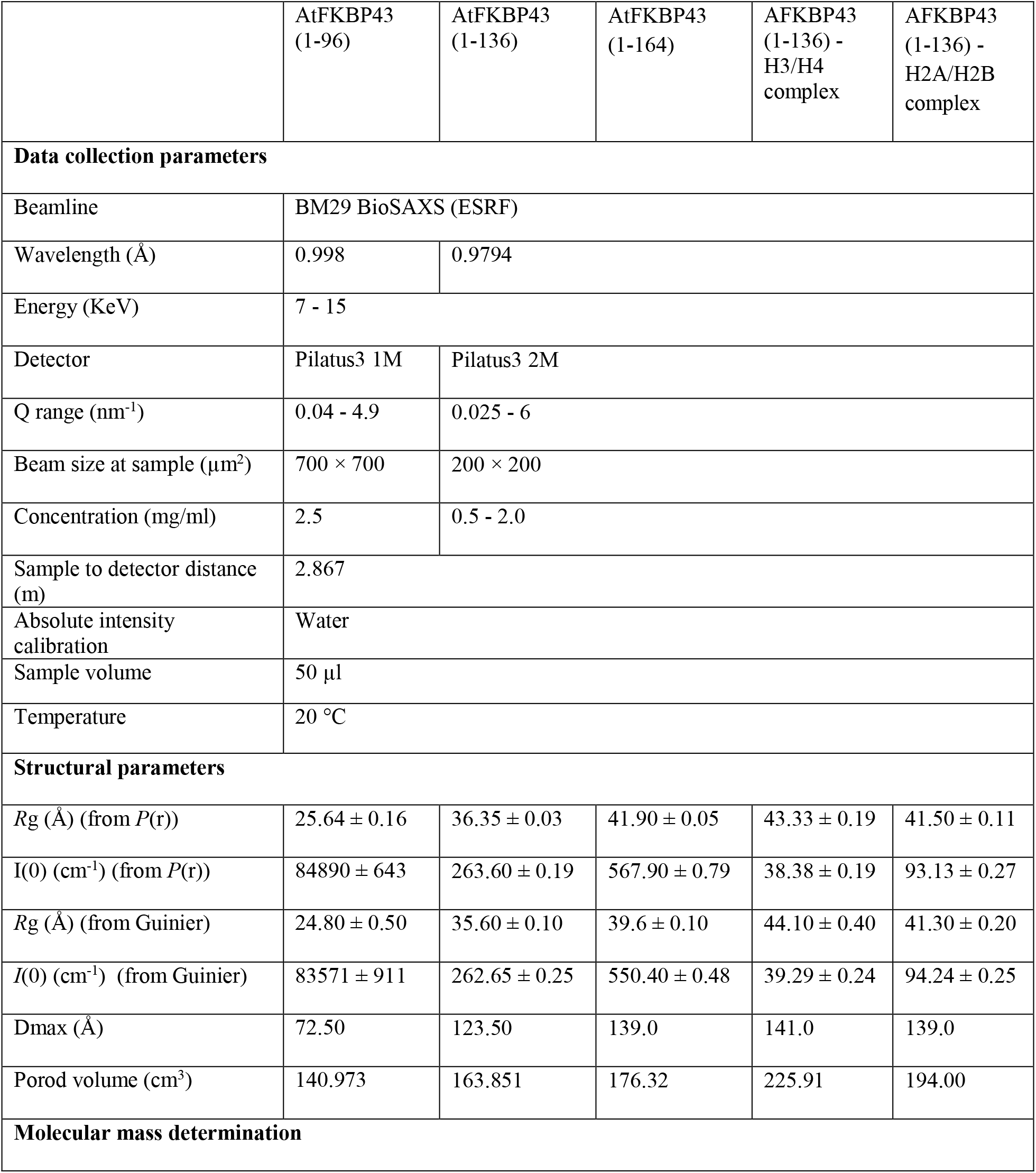
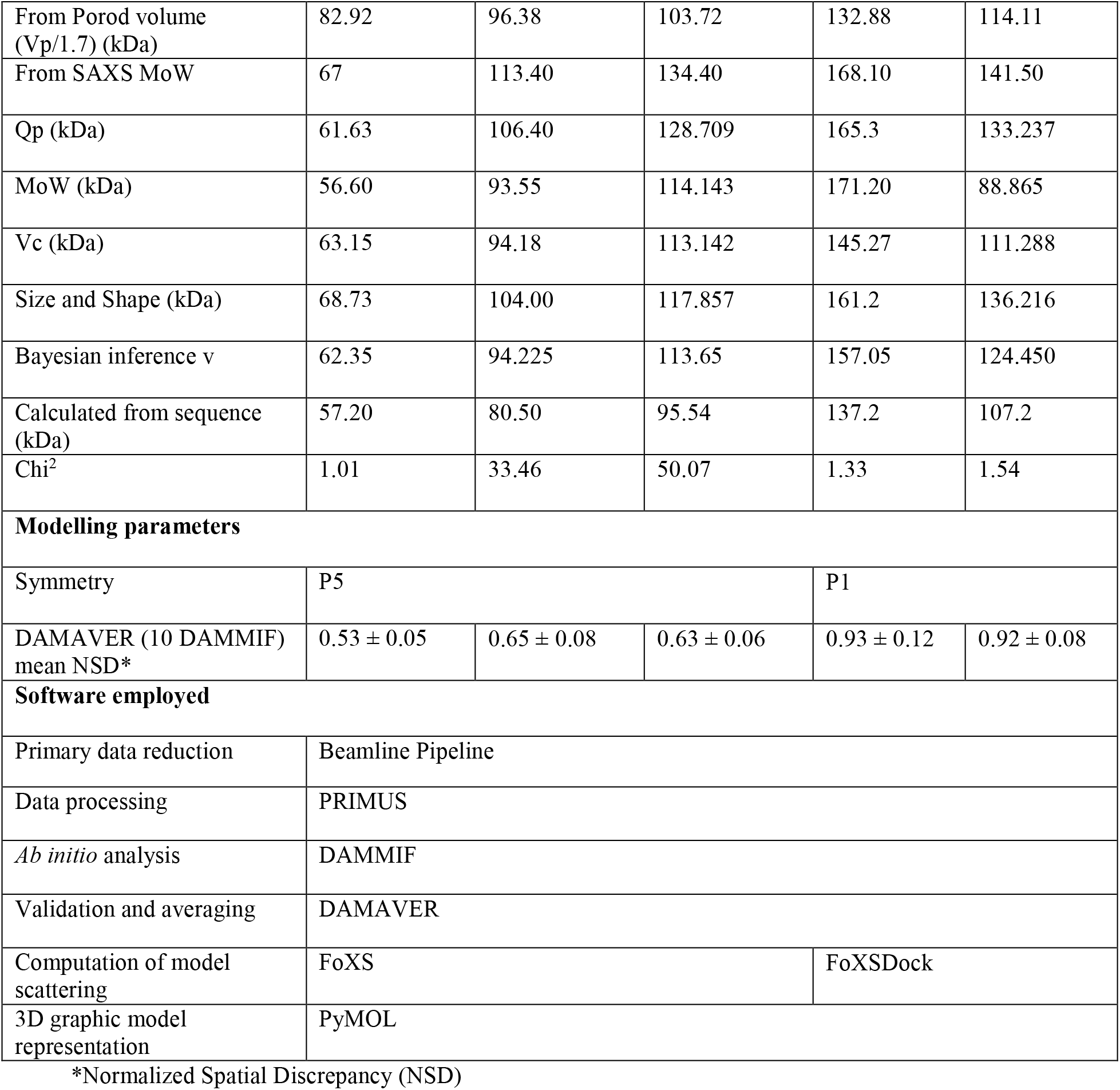
SAXS data collection and structure parameters for AtFKBP43 NTDs and their histone complexes.

The molecular mass estimated using Porod volume (*Vp*/1.7) from ATSAS (36) was observed to be 82 kDa, 96 kDa, and 103 kDa for AtFKBP43 (1-96), (1-136), and (1-164), respectively. The overlaid theoretical scattering against experimental scattering for AtFKBP43 (1-96) displayed a significant χ^2^ (goodness of fit) value of 1.01 whereas AtFKP43 (1-136) and (1-164) yielded high χ^2^ values (33.46 and 50.07 respectively) suggesting an elongated nature attributable to the conformational flexibility of the disordered C-terminal acidic stretches. The overall SAXS structural parameters suggest that AtFKBP NTDs exist as pentamer in solution.

### The crystal structure of AtFKBP43 NTD reveals a characteristic nucleoplasmin fold

The N-terminal core domain of AtFKBP43, spanning residues 1-96, did not yield any crystals. However, a slightly longer stretch of AtFKBP43 NTD (spanning residues 1-110) yielded crystals that diffracted up to 2.3 Å. Molecular replacement was used to solve the structure, using the structural coordinates of the AtFKBP53 NTD monomer (PDB id: 6J2Z) (29) as a search model. The solved structure had four individual pentamers without any decamer-like conformation. The statistics for crystal data collection, data processing, and subsequent refinement can be seen in **Table 3**.

**Table 3:** AtFKBP43 (1-110) crystal data collection, processing, and refinement statistics.

| Parameter | AtFKBP43 (1-110) |
| --- | --- |
| **Data collection and processing** |  |
| Beamline | ID23-1, Grenoble, France |
| Detector type | Dectris Eiger X 4M |
| Wavelength (Å) | 0.9794 |
| Data collection temperature (K) | 100 |
| Space group | P2<sub>1</sub> |
| a, b, c (Å) | 53.06, 203.91, 94.55 |
| $\alpha$, $\beta$, $\gamma$ (°) | 90.00, 96.14, 90.00 |
| Resolution (Å) | 48.28-2.29 (2.42-2.29) |
| R<sub>merge</sub> (%) | 12.9 (67.8) |
| I/σI | 10.4 (2.1) |
| **Refinement** |  |
| CC (1/2) (%) | 0.99 (0.89) |
| No. of unique reflections | 88501 (8417) |
| Completeness (%) | 100.0 (99.3) |
| Multiplicity | 6.6 (6.5) |
| Wilson B-factor (Å<sup>2</sup>) | 47.0 |
| Solvent content (%) | 44.01 |
| No. of molecules in ASU | 20 |
| No. of unique reflections | 88106 |
| R<sub>work</sub>/R<sub>free</sub> (%) | 22.3/25.8 |
| Total no. of non-H atoms | 13402 |
| No. of water molecules | 10 |
| Mean B-factor (Å<sup>2</sup>) | 67.0 |
| R.m.s. deviations: |  |
| Bond lengths (Å) | 0.004 |
| Bond angles (°) | 0.81 |
| Ramachandran plot values (%): |  |
| Favoured / Outliers | 100/ 0 |

One out of the four pentamers, that had improved electron density was used to analyze the structural details and to prepare the figures. Each monomeric unit had electron density only up to the 97^th^ residue suggesting that the C-terminus, comprised of up to 110 residues and His-tag is unstructured. The N and C termini of all five monomers of a pentamer go into and come out of the same proximal face, similar to AtFKBP53 NTD, an organization typical of nucleoplasmins **(Fig. 3A)**.

**Figure 3.**
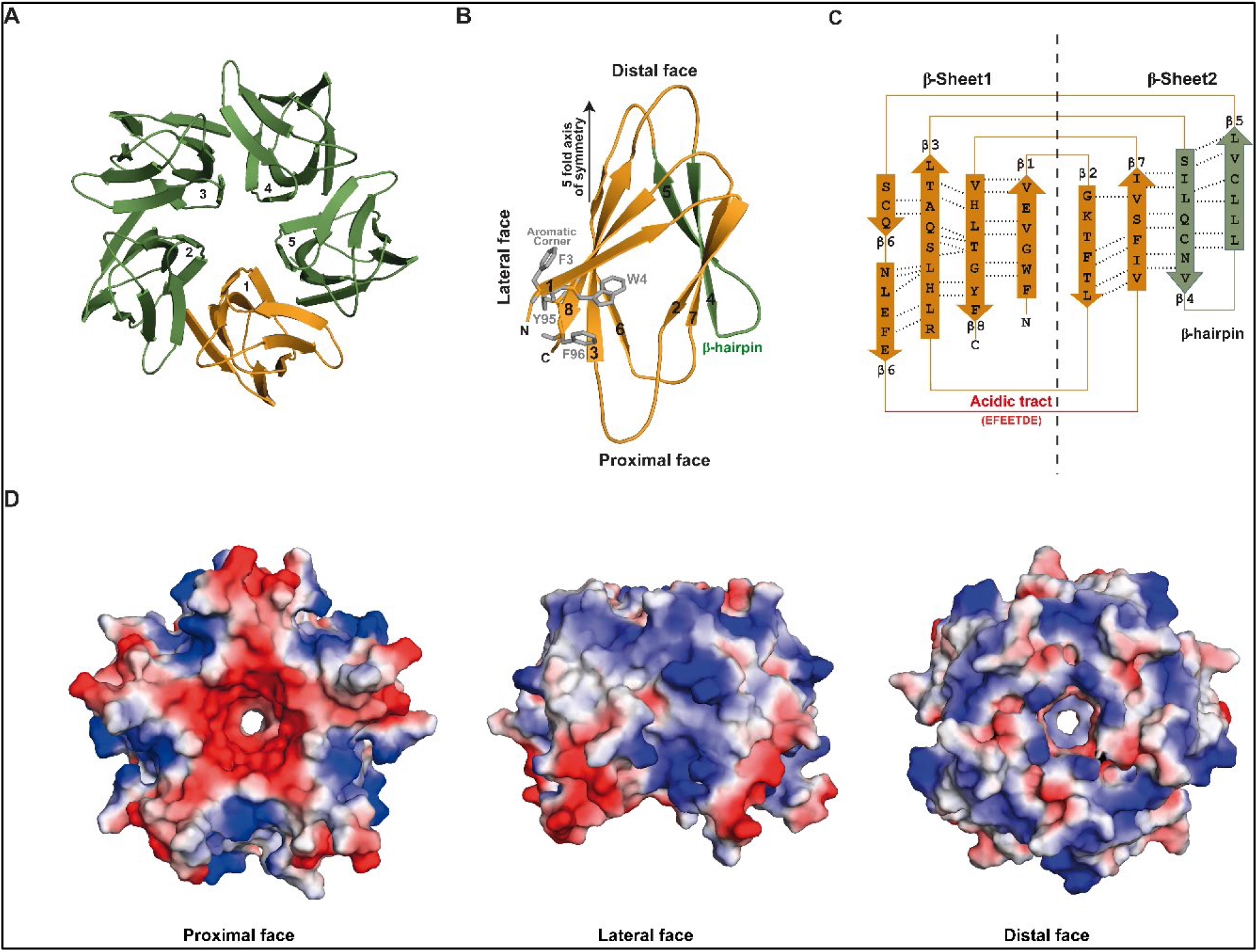
Crystal structure of AtFKBP43 NTD. **(A)** Cartoon representation of the AtFKBP43 NTD pentamer crystal structure (green), wherein one monomer is highlighted in orange. The figure shows the view of the distal face. **(B)** Cartoon representation of AtFKBP43 NTD monomer structure having a conserved tear-drop shape; the distal, lateral, and proximal faces are labelled and the five-fold axis of symmetry is shown with a black arrow. The N-terminal end goes in and the C-terminal ends comes out of the proximal face. The A1 acidic tract (red), the aromatic corner made up of Phe3, Trp4, Tyr95, Phe96 (stick representation, with side-chains in grey), and the β-hairpin motif made out of β4 and β5 (in green) are also shown. **(C)** 2D jelly-roll topology diagram of the AtFKBP43 NTD monomer, displaying the arrangement and the hydrogen bonding patterns of the β-strands with the latter marked in dotted lines. The acidic tract A1 made up of residues EETDE and the β-hairpin motif are also labelled. The dotted black line indicates the two β-sheet halves - β-sheet 1 and β-sheet 2, representing the sandwich-type structural organization within a monomer. **(D)** Surface charge distribution of AtFKBP43 NTD. The electrostatic surface features of AtFKBP43 NTD in its proximal, distal, and lateral faces. PyMOL has been used to generate the surface potential that is coloured in a linear fashion from red (-kT) to blue (+kT).

A monomer of AtFKBP43 NTD is defined by a signature beta-sandwich fold of the nucleoplasmin family of proteins, consisting of eight β-strands folded in a jelly roll topology, thus confirming it to be a nucleoplasmin, like AtFKBP53. Each individual β-strand runs parallel to the 5-fold axis of the pentamer **(Fig. 3B)**. Out of the eight β-strands, the β-hairpin made up of β4 and β5 twists out and remains farthest from the fivefold axis while β6 lies closest to the axis. Hydrogen bonds between the antiparallel β-strands stabilize the jelly roll topology of AtFKBP43 NTD **(Fig. 3C)**.

The asymmetric distribution of four bulky aromatic residues on strand β1 (Phe3 and Trp4) and β8 (Tyr95 and Phe96), and the conserved β-hairpin motif comprised of β4 and β5 strands provide each monomer a tear-drop shape **(Fig. 3B)**. Within a pentamer, the β-hairpin motif of one monomer is located at the interface between the two monomers and stays perpendicular to the β-sheet of the adjacent monomer.

### AtFKBP43 NTD structure compares well with other nucleoplasmins

The crystal structure of the AtFKBP43 NTD monomer was aligned with the structures of the evolutionarily closest nucleoplasmins such as AtFKBP53 NTD, DmNLP, and HsNPM1, and the topology compared. The NTD of AtFKBP43 has a similar β-barrel topology, with an average RMSD value of 1.6 Å (for Cα atoms) with these structures (29, 34, 37) **(Fig. S3A)**. Although the beta-barrel topology is identical, a significant difference in the length and the trajectory of the β-hairpin motif of AtFKBP43 NTD with HsNPM1 and DmNLP could be observed. The length of β5, which forms part of the β-hairpin motif in AtFKBP43, is 15.0 Å in comparison to 24.1 Å in HsNPM1. The β-hairpin motif in DmNLP is packed against the lateral surface in contrast to the extended conformation in AtFKBP43 NTD, with a 9.5 Å displacement from the lateral surface **(Fig. S3B)**.

AtFKBP43 NTD has a flexible nine residue-long loop between β2 and β3 strands instead of a 3_10_ helix in the AtFKBP53 NTD structure **(Fig. S3C)**. Unlike, the crystal structures of XlNPM and XlNO38-NPM, in AtFKBP43 the conserved AKDE motif, GSGP motif and K-loop have an ATNR motif, GPRS motif and Thr61, respectively **(Fig. S3D)**.

Additionally, the A1 tract is present between β6 and β7 for AtFKBP43 NTD and AtFKBP53 NTD, whereas it is present between β2 and β3 for other nucleoplasmins. In AtFKBP43 NTD the discontinuous, acidic patch comprised of four acidic residues (EETDE) on the proximal face is primarily restricted around the central pore, whereas the A1 tract of AtFKBP53 NTD gives a continuous acidic patch (DDDDE) to its entire proximal face. The central cavity of the AtFKBP43 NTD also has an acidic charge. The lateral and distal face surface of AtFKBP43 NTD appears predominantly basic in charge distribution **(Fig. 3D)**.

### AtFKBP43 NTD pentamer is highly stable

Nucleoplasmin core domains are highly stable against chemical and physical challenges, which is true for AtFKBP53 NTD as well (29). Analytical-SEC experiments to evaluate the stability of AtFKBP43 NTD (1-96) pentamer under thermal and chemical unfolding conditions indicated the protein to be stable up to 60 °C, 2 M sodium chloride, and 4 M urea. No shift in elution volume and no appearance of new peaks until 80 °C, though with decreasing peak heights, suggests that the pentamer does not fall apart into lower oligomeric or monomeric forms **(Fig. 4A)**. An increase in urea concentration above 4 M resulted in a delay in the elution peak, suggesting that the pentameric protein fell apart into lower oligomeric forms **(Fig. 4, B and C)**.

**Figure 4.**
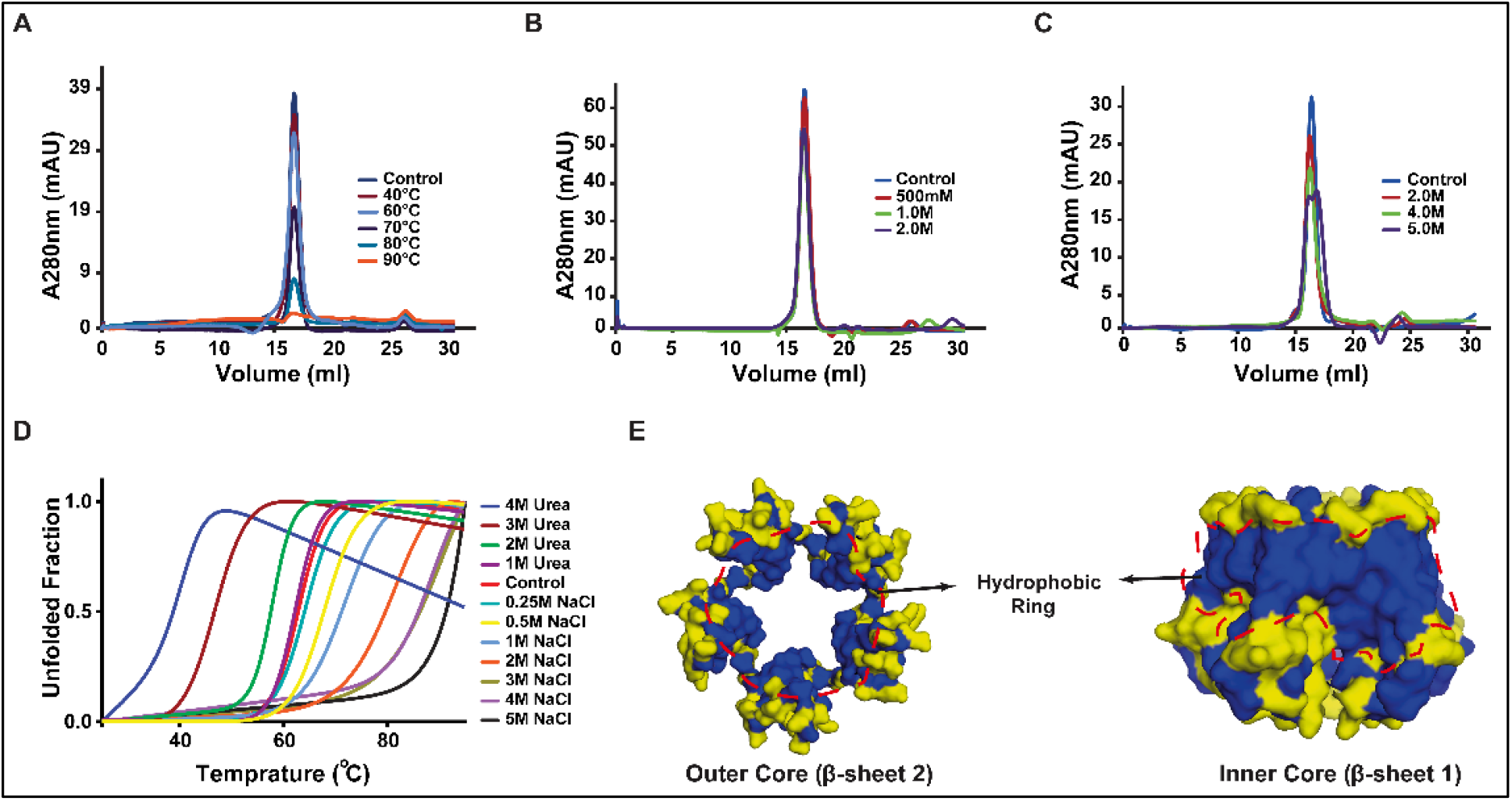
Stability of AtFKBP43 NTD pentamer. **(A)** Analytical size-exclusion chromatogram of AtFKBP43 NTD (1-96) pentamer subjected to different temperatures: 26 ºC (control; blue), 50 ºC (brown), 60 ºC (light green), 70 ºC (purple) and 80 ºC (light blue), showing that the protein is heat stable up to 60 °C. **(B)** Analytical size-exclusion chromatogram of AtFKBP43 NTD pentamer at increasing NaCl concentrations: 150 mM (blue), 300 mM (black), 600 mM (orange) and 1.0 M (red) showing that the pentameric organization is quite salt-resistant. **(C)** Analytical size-exclusion chromatogram of AtFKBP43 NTD pentamer under increasing urea concentrations: in absence of urea (blue), with 2.0 M urea (brown), with 4.0 M urea (light green), and with 6.0 M urea (purple), showing stability until about 4.0 M urea. **(D)** Thermal denaturation profile of AKBP43 NTD pentamer monitored by circular dichroism at 222 nm. The protein was treated with varying concentrations of urea and NaCl. **(E)** A surface representation of the different orientations of the beta-sheets making up the outer and inner layers of the AtFKBP43 NTD pentamer displaying the apolar and hydrophobic residue distribution. The apolar and hydrophobic residues are coloured in blue and the rest are in yellow. They are seen to form a concentric ring-like network, thus stabilizing the pentamer structure.

Further, thermal ramping Circular Dichroism (CD) experiments with AtFKBP43 NTD (1-96) alone and in the presence of increasing concentrations of either NaCl or urea indicated a melting temperature (Tm) of 63.31 °C that concomitantly increased with increasing salt to 107.35 °C at 5 M NaCl while decreasing to 42 °C at 4 M urea. The Tm of AtFKBP43 NTD (1-96) corroborated well with the analytical-SEC results **(Fig. 4D and Table S2)**.

A closer observation indicated that the stability of the protein under harsh chemical and temperature conditions might be attributed to the similarity in structural elements and distribution of hydrophobic and apolar residues **(Fig. S4A)** (20, 29, 34, 37-39). In AtFKBP43 NTD, the inner and outer layers of the pentamer core made out of beta sheet-1 and beta sheet-2 of all the monomers, respectively, predominantly contain hydrophobic and apolar residues, side chains of which interact to form a contiguous ring, thus adding stability to the structure **(Fig. 4E)**. Similar interactions exist between the highly thermostable archaeal chaperonin subunits called thermosome, forming a hydrophobic ring (40)

Moreover, reports suggest that the N-terminal core domain of nucleoplasmins exhibits resistance to protease, whereas the C-terminal flexible tail region displays protease sensitivity (20, 41) The proteinase K-treated AtFKBP43 (1-96), (1-136), and (1-164) samples showed elution volumes in analytical SEC similar to untreated AtFKBP43 (1-96) pentamer, indicating that AtFKBP43 NTD is resistant to cleavage but suggested that the C-terminal unstructured acidic tail region gets cleaved into smaller peptides **(Fig. S4B)**. SDS-PAGE analysis of the eluted fractions also supported this observation **(Fig. S4C)**.

### AtFKBP43 NTD with its C-terminal arms, but not the NTD alone interacts specifically with histone H2A/H2B and H3/H4

Nucleoplasmins are reported to interact with H2A/H2B and/or H3/H4 histone oligomers (20, 39, 42-44). AtFKBP53 NTD also interacts with H2A/H2B as well as H3/H4 oligomers (29). Moreover, previous reports suggest that the disordered acidic stretches significantly enhance histone affinity of the nucleoplasmin core domain, which is inevitable for histone chaperoning activity (44). Hence, the interaction of AtFKBP43 (1-96), (1-136), and (1-164) and the role of its acidic stretches in regulating histone interaction were explored. In addition, to determine the minimum number of acidic residues required for histone interaction, AtFKBP43 NTD containing half of the A2 acidic tract, i.e., spanning residues 1-110, was also prepared.

The interaction of AtFKBP43 NTD with H2A/H2B and H3/H4 oligomers examined using Ni-NTA pull-down assays suggested that AtFKBP43 (1-136) and (1-164) interact with H2A/H2B and H3/H4 oligomers. AtFKBP43 (1-110) pulled down H3/H4 to a much lesser extent, but not H2A/H2B, while AtFKBP43 (1-96) did not interact with the histone oligomers **(Fig. S5, A and B)**.

To validate the results from the pull-down assays, analytical-SEC experiments were performed. It was found that AtFKBP43 (1-136) and AtFKBP43 (1-164) indeed interact and form complexes with H2A/H2B and H3/H4, independently as evident from the shift observed in the elution volume of the complexes in analytical-SEC as well as the subsequent peak fraction analyses on SDS-PAGE gel **(Fig. 5, A and B)**. Next, the binding stoichiometry of AtFKBP43 (1-136) and (1-164) with histone oligomers were analyzed using SV-AUC experiments. This gave sedimentation coefficient values and molecular mass estimates for AtFKBP43 (1-136) and AtFKBP43 (1-164) pentamers in complex with either H2A/H2B dimer or H3/H4 tetramer, and those corresponded to complex formation revealing a 1:1 binding stoichiometry in each case **(Fig. 5, C and D; Fig. S5, C-H)**. A decrease in the Stokes radius, frictional ratio, and Smax/S values **(Table 1)** suggested that AtFKBP43 histone oligomers complexes adopt a globular conformation.

**Figure 5.**
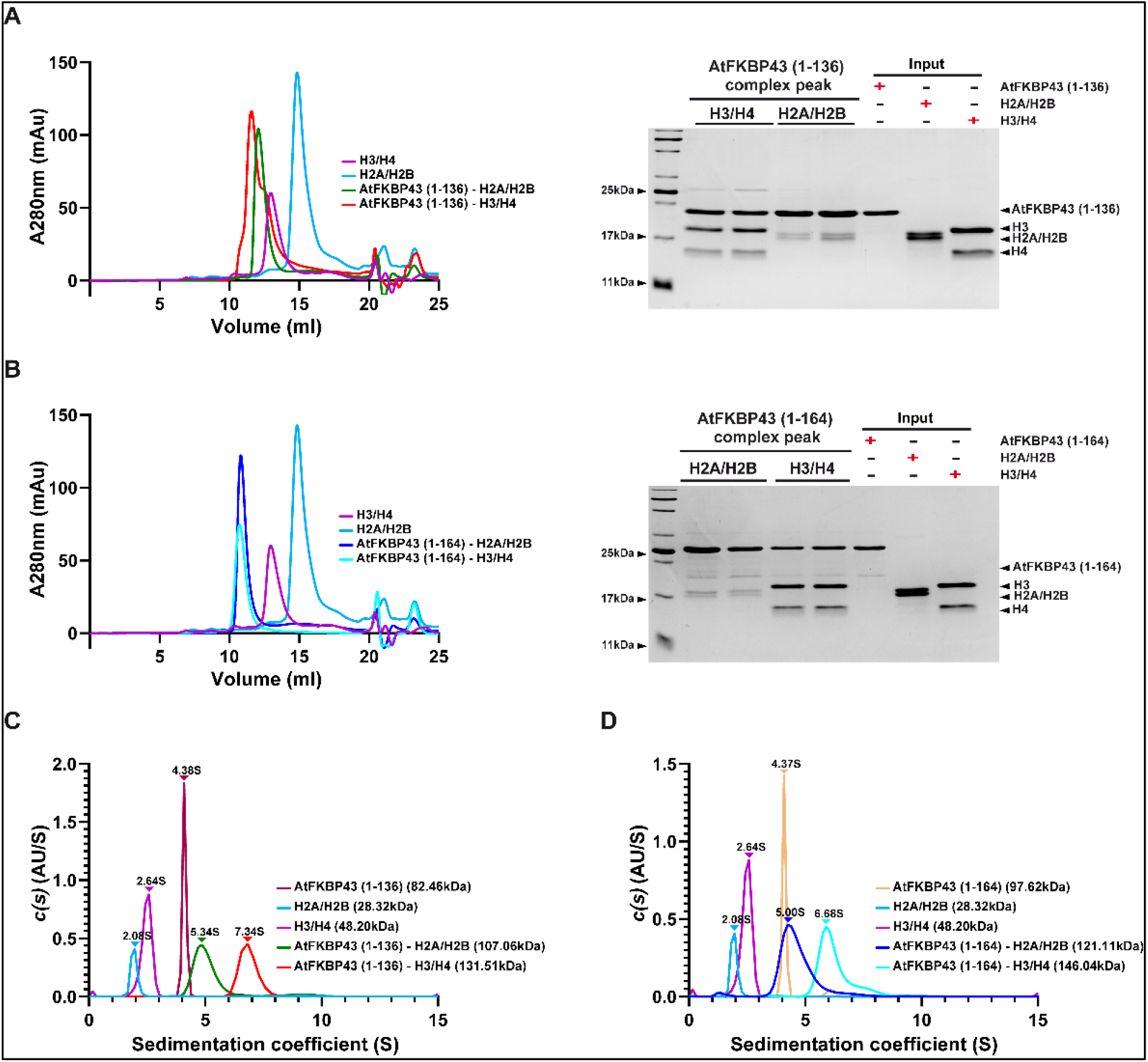
Analytical-SEC and SV-AUC analysis of AtFKBP43 (1-136) and (1-164) pentamers and their histone oligomer complexes. **(A)** Analytical SEC profiles of H2A/H2B (light blue), H3/H4 (pink), AtFKBP43 (1-136)-H2A/H2B mixture (green), and AtFKBP43 (1-136)-H3/H4 mixture (red) indicating the formation of stable complexes. Peak fractions from analytical-SEC run on 18% SDS-PAGE gel showing complex formation (right panel). **(B)** Analytical SEC profiles of H2A/H2B (light blue), H3/H4 (pink), AtFKBP43 (1-164)-H2A/H2B mixture (blue), and AtFKBP43 (1-164)-H3/H4 mixture (cyan) indicating the formation of stable complexes. 18% SDS-PAGE gel image showing the peak fractions obtained from the SEC, confirming complex formation (right panel). **(C)** Comparison of sedimentation coefficient (S) to c(S) distribution plot for AtFKBP43 (1-136) (dark brown), AtFKBP43 (1-136) – H2A/H2B complex (green), AtFKBP43 (1-136) – H3/H4 complex (red), H3/H4 (pink) and H2A/H2B (light blue). The sedimentation coefficient (S20, w) and molecular mass for each protein sample determined using the SEDFIT software (in parentheses) are also shown. **(D)** Comparison of sedimentation coefficient (S) to c(S) distribution plot for AtFKBP43 (1-164) (orange), AtFKBP43 (1-164) – H2A/H2B complex (blue), AtFKBP43 (1-164) – H3/H4 complex (cyan) H3/H4 (pink) and H2A/H2B (light blue). The sedimentation coefficient (S20, w) and molecular mass for each protein sample determined using the SEDFIT software (in parentheses) are also shown.

Further, the overall envelope structures of AtFKBP43 in complex with H2A/H2B and H3/H4 were obtained using SAXS experiments. Since our pull-down and analytical-SEC experiments revealed the importance of the A1 stretch in interaction with histone oligomers, AtFKBP43 (1-136) was chosen for SAXS studies. Guinier plot **(Fig. S6B)**, Kratky plot **(Fig. S6C)**, paired distance distribution **(Fig. S6D)** suggested that AtFKBP43 (1-136)-H2A/H2B and AtFKBP43 (1-136)-H3/H4 complexes are homogenous, folded and adopt globular conformation. The Dmax and Rg values were estimated using GNOM and additional structural parameters for AtFKBP43 (1-136)-H2A/H2B and AtFKBP43 (1-136)-H3/H4 are shown in **Table 2**. The molecular masses for AtFKBP43 (1-136)-H2A/H2B and AtFKBP43 (1-136)-H3/H4 complexes estimated using the Porod volume (*Vp*/1.7) from ATSAS (36) were found to be 114.11 kDa and 132.88 kDa, respectively, suggesting a 1:1 stoichiometry thus in agreement with the results from SV-AUC. The docked models for AtFKBP43 (1-136) in complex with H2A/H2B and H3/H4 gave χ^2^ values of 1.82 and 1.60, respectively, superimposing quite well into their respective bead models **(Fig. 6, A and B)**.

**Figure 6.**
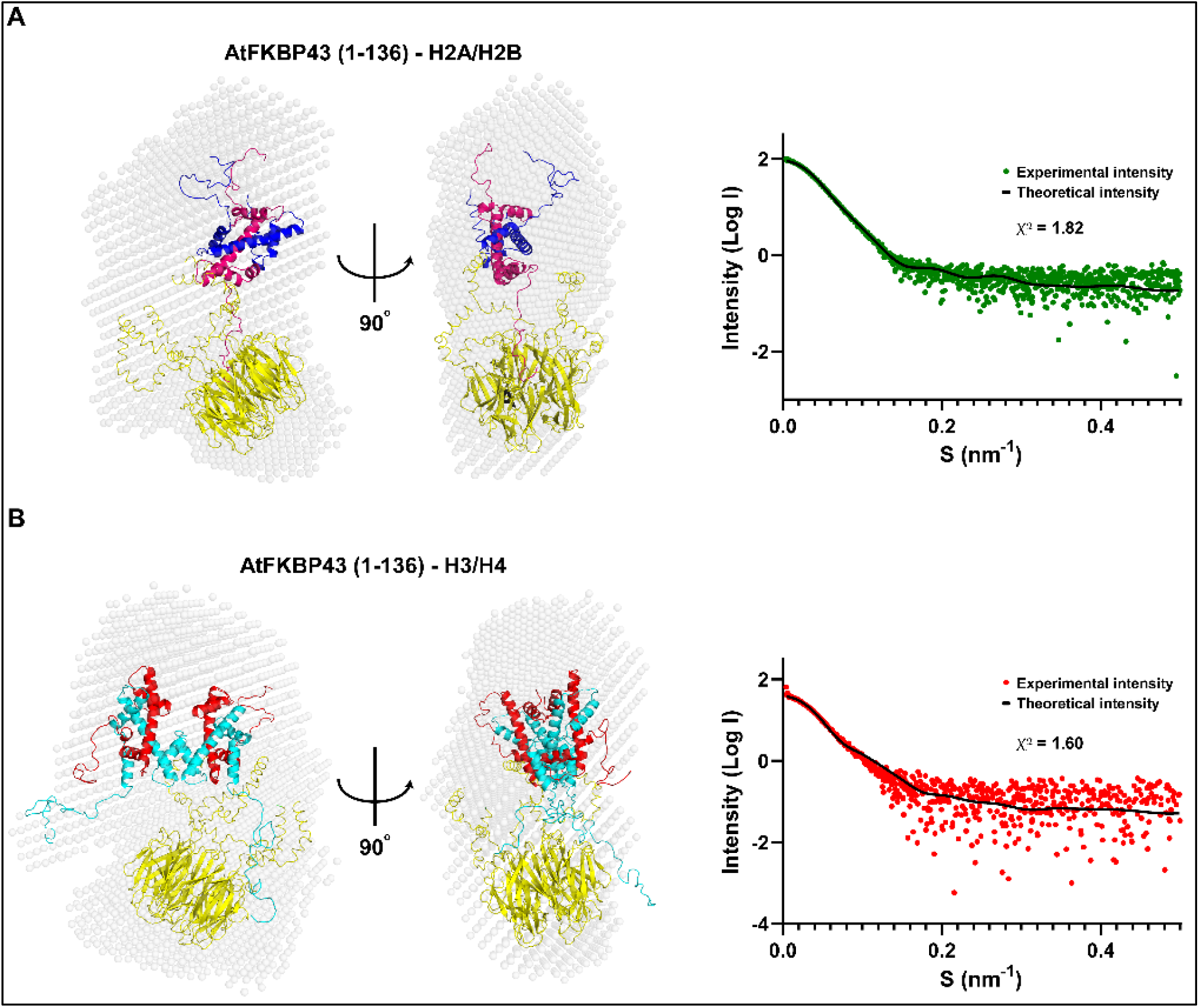
SAXS derived *ab initio* envelope and model structures of AtFKBP43 (1-136) in complex with H2A/H2B and H3/H4. **(A)** SAXS-based low-resolution envelope (spheres denoting the dummy residues) of the AtFKBP43 (1-136)-H2A/H2B complex with the fit to best rigid body model of AtFKBP43 (1-136)-H2A/H2B calculated from FoXSDock. The structure model of AtFKBP43 (1-136) is represented in yellow, H2A is represented in hot pink, and H2B is in blue. The scattering curve from the experimental SAXS profile of AtFKBP43 (1-136)-H2A/H2B (green dots) overlaid with theoretical scattering curve from the rigid body model (black line) calculated from FoXSDock, with the corresponding χ^2^ value is presented (right panel). **(B)** SAXS-based low-resolution envelope of the AtFKBP43 (1-136)-H3/H4 complex with the fit to best rigid body model of AtFKBP43 (1-136)-H3/H4 calculated from FoXSDock. The structure model of AtFKBP43 (1-136) is represented in yellow, H3 is represented in cyan, and H4 is in red. The scattering curve from the experimental SAXS profile of AtFKBP43 (1-136)-H3/H4 (red dots) overlaid with the theoretical scattering curve from the rigid body model (black line) calculated from FoXSDock, with the corresponding χ^2^ vale is presented (right panel).

To investigate the binding preference of AtFKBP43 (1-136) and AtFKBP43 (1-164) for H2A/H2B and H3/H4 histone oligomers, ITC experiments were performed. Upon titrating AtFKBP43 (1-136) and AtFKBP43 (1-164) against H2A/H2B dimer, separately, the reactions were observed to be endothermic and entropically driven with a K_d_ of 0.46 **±** 0.03 µM for AtFKBP43 (1-136), and 0.06**±** 0.006 µM for AtFKBP43 (1-164). The calculated thermodynamic parameters are also presented **(Fig. 7A and Fig. S7A)**.

**Figure 7.**
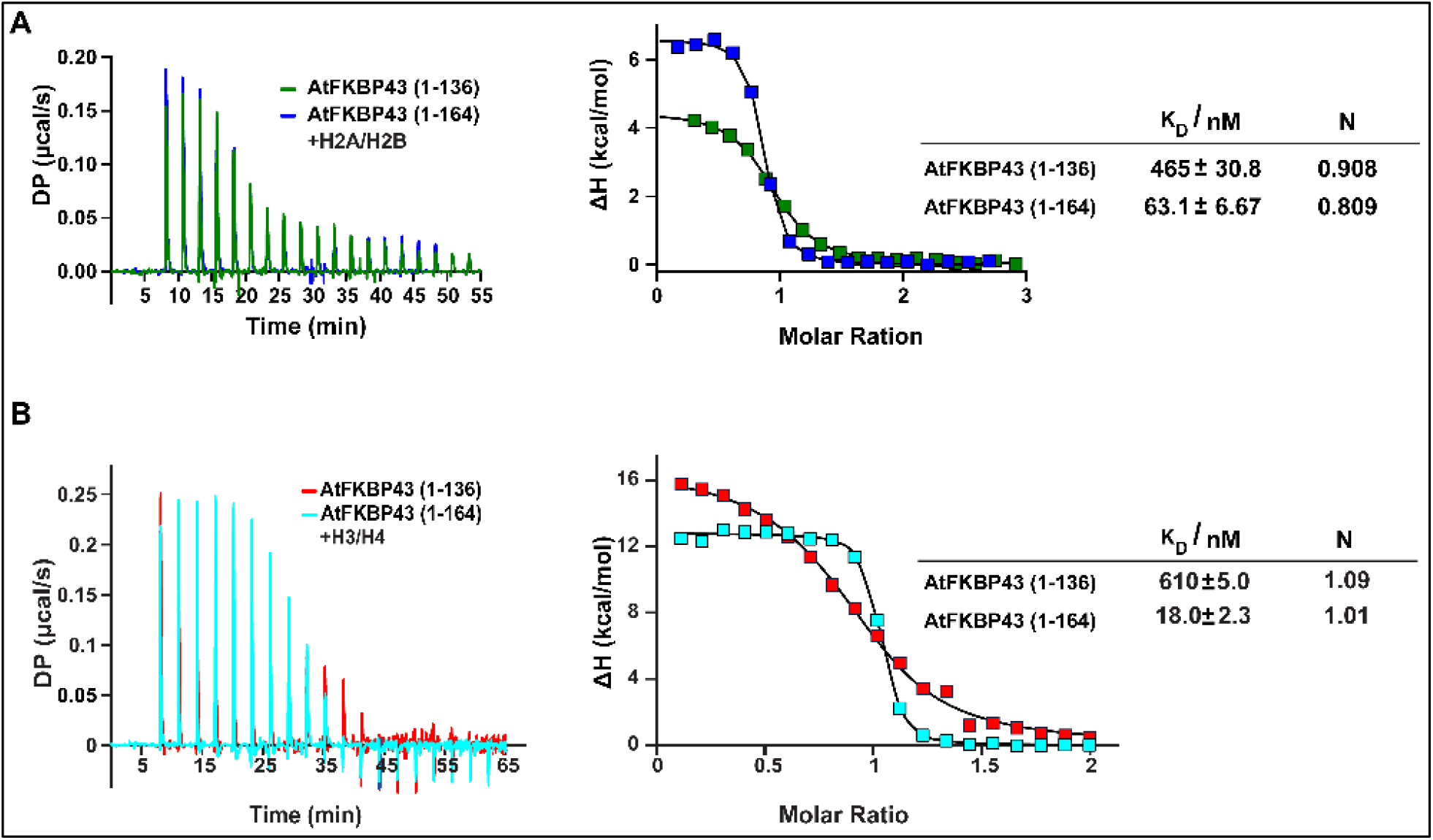
Isothermal titration calorimetry of AtFKBP43 (1-136) and AtFKBP43 (1-164) with histone oligomers. Titrations of AtFKBP43 (1-136) and AtFKBP43 (1-164) against H2A/H2B and H3/H4 separately, were performed by ITC at 25 °C. **(A)** The left panel shows the heat changes occurring upon injections of either AtFKBP43 (1-136) (in green) or AtFKBP43 (1-164) (in blue) to H2A/H2B. The right panel presents the isotherms obtained upon fitting the ITC data to a ‘one set of site’ binding model. The right panel shows the Kd and stoichiometry values. **(B)** The left panel shows the heat changes occurring upon injections of eitherAtFKBP43 (1-136) (in red) or AtFKBP43 (1-164) (in light blue) to H3/H4. The right panel presents the isotherms obtained upon fitting the ITC data to a ‘one set of site’ binding model. The right panel shows the Kd and stoichiometry values.

Similarly, upon titrating AtFKBP43 (1-136) and AtFKBP43 (1-164) against H3/H4 tetramer, separately, endothermic reactions were observed with a K_d_ value of 0.61 **±** 0.04 µM for AtFKBP43 (1-136) and 0.018 **±** 0.002 µM for AtFKBP43 (1-164). The calculated thermodynamic parameters are presented as well **(Fig. 7B and Fig. S7B)**. Both the interactions showed a 1:1 binding stoichiometry, thus being in agreement with the results from SV-AUC and SAXS experiments.

The interpretation of the ITC data showed that hydrophobic interactions contribute significantly to the stability of the histone oligomer-AtFKBP43 complex, as reported earlier for other nucleoplasmins (44, 46). However, the abrogation of complex formation upon deletion of the acidic tract in AtFKBP43 reflected that electrostatic interaction is critical for complex formation. A pull-down assay further corroborated this, wherein the His-tagged AtFKBP43 (1-136) was incubated separately with untagged H2A/H2B **(Fig. S8A, left panel)** and H3/H4 **(Fig. S8A, right panel)**. Similarly, His-tagged AtFKBP43 (1-164) was incubated separately with H2A/H2B **(Fig. S8B, left panel)** and H3/H4 **(Fig. S8B, right panel)** with concomitantly increasing salt concentration conditions. Elution profiles showed that the H3/H4 complex was stable up to 1.0 M NaCl concentration, while the H2A/H2B complex appeared to be stable only up to 0.3 M NaCl. These observations imply that the hydrophobic interactions are significant in providing stability to the H3/H4-AtFKBP43 complex, but to a much lesser extent for the H2A/H2B-AtFKBP43 complex.

## DISCUSSION

A major step in deciphering the mechanism of chromatin remodelling and its effect on gene regulation is identifying the nuclear factors involved in the nucleosome assembly and disassembly process. It has already been reported that histone chaperones play a significant role in regulating chromatin dynamics. However, to date, the functional mechanisms of only a few histone chaperones have been characterized, and many more remain to be uncovered. Like AtFKBP53, AtFKBP43 is also a multi-domain FKBP from *A. thaliana*, and herein, it has been shown to have a nucleoplasmin domain at its N-terminus and interacts with H2A/H2B and H3/H4 oligomers, thereby suggesting its role as a histone chaperone. However, unlike AtFKBP53, where the core domain could interact with the histone oligomers, AtFKBP43 binds to the histone oligomers through its highly acidic charged extensions protruding from the core domain.

Eight structures have been reported from the family of nucleoplasmins to date, including XlNPM1, XlNPM2, HsNPM1, HsNPM2, MmNPM1, DmNLP DmFKBP39 NLP, and AtFKBP53 NTD. All these structures have shown a conserved homo-pentameric conformation. Here, the first structural report of AtFKBP43 NTD shows the domain to have a pentameric nucleoplasmin fold, with the monomer possessing a tear-drop shape.

Various oligomeric states of nucleoplasmins have been suggested owing to possible repulsive forces between the subunits generated due to the highly acidic charged or phosphorylated tail regions (47). As the AtFKBP43 constructs were all recombinantly expressed in bacteria, only the effect of acidic stretches could be tested, and not the effect of phosphorylation states. The proteins having A2 and A3 stretches were still pentameric in their organization, though the C-terminal flexible acidic stretches protruding out from the proximal face contribute to their overall elongated conformation in contrast to the compact globular nature of the AtFKBP43 NTD (1-96) core domain. Unlike nucleoplasmins such as XlNPM and XlNO38-NPM, the absence of AKDE/GSGP motif and the K-loop in the AtFKBP43NTD structure, could be a reason for not taking up a stable decameric conformation.

The pentameric states of XlNPM2, HsNPM2, DmNLP, and AtFKBP53 NTD have been reported to be resistant to high temperatures. Further examination of their nucleoplasmin domains has shown that the stability to extreme denaturing conditions of the pentamer is provided by the hydrophobic interactions between the inner and the outer core layers of the nucleoplasmin pentamer (20, 29, 34, 37-39) which hold for AtFKBP43 NTD as well. Moreover, AtFKBP43 (1-96) pentamer shows protease resistance in agreement with the previous reports on other nucleoplasmins (20, 41).

In contrast to AtFKBP53 NTD, AtFKBP43 NTD (1-96) did not interact with histone oligomers. The discontinuous A1 acidic tract in AtFKBP43 NTD (1-96), in contrast to a continuous one in AtFKBP53 NTD, could be a reason for the former not interacting with the histones. Like HsNPM2, AtFKBP43 requires the longer acidic tract A2 [AtFKBP43 (1-136)] to interact stably with the histone oligomers H3/H4 and H2A/H2B.

Electron microscopy and SAXS studies have displayed that one pentamer of XINPM2 can accommodate five H2A/H2B dimers whereas two pentamers of XINPM2 are required to accommodate one H3/H4 tetramer and histone octamer, individually on its proximal face (32, 46). However, the stoichiometry of the interaction of AtFKBP43 (1-136) pentamer with the H2A/H2B and H3/H4 histone oligomers is 1:1, which is true for AtFKBP53 NTD as well (29). This implies that FKBP nucleoplasmins might possess a different mode of association for histone oligomers as compared to the canonical nucleoplasmins. Further, SAXS and AUC data demonstrate the AtFKBP43 nucleoplasmin-histone oligomer complexes to adopt a globular conformation in contrast to the elongated nature of the AtFKBP43 nucleoplasmin core with the C-terminal acidic stretches alone. The globular shape may be attributed to stabilizing the otherwise flexible C-terminal tail of AtFKBP43 (1-136) and the N-terminal tails of histone oligomers upon interaction, as has been reported for XINPM2-H2A/H2B complex (32). The binding affinity for both the histone oligomers was observed to be lower than what has been reported for recombinant XlNPM2 containing the A2 acidic tract (42). The low affinity could be due to the differences in the distribution of acidic residues within the A2 tract of the proteins. In AtFKBP43, the A2 tract has 13 acidic residues mixed with 17 polar and hydrophobic residues. On the other hand, in XlNPM2, the A2 tract has 19 acidic residues, interspersed with only seven non-charged, hydrophobic and polar residues put together. Further, in AtFKBP43, the inclusion of the acidic tract A3 increases its binding affinity to the histone oligomers.

There are reports for recombinant XlNPM2 core domain interacting with H2A/H2B dimer (32) and AtFKBP53 NTD interacting with H2A/H2B and H3/H4 (29), which shows that the interaction is favoured majorly through hydrophobic interactions. However, electrostatic and polar interactions also play an important role, either by orienting the binding partners properly or aiding direct binding through charge compensation (48-50). Our ITC and salt gradient pull-down assay of H2A/H2B and H3/H4 histone oligomers with AtFKBP43 (1-136) and AtFKBP43 (1-164) indicate that though electrostatic interactions drive the association, the binding is probably stabilized through hydrophobic interactions. However, hydrophobic interactions appear less predominant for H2A/H2B than H3/H4 for complex formation with AtFKBP43, similar to the AtFKBP53 core nucleoplasmin domain (29). The binding affinity of AtFKBP53 NTD to H2A/H2B is between those for the AtFKBP43 (1-136) and AtFKBP43 (1-164). However, the binding affinity of AtFKBP53 NTD to H3/H4 is lower than both AtFKBP43 (1-136) and AtFKBP43 (1-164).

In conclusion, our results presented here, put forward a notion that FKBP nucleoplasmins might have evolved a distinct mode of histone oligomers recognition, contrary to classical nucleoplasmins, that govern their stoichiometry of interaction. The structural basis of these interactions needs further investigation. Though unlike AtFKBP53 NTD, the acidic stretches on the tail region for AtFKBP43 NTD are required for histone interaction, a previous study has found that the individual null mutants of AtFKBP53 and AtFKBP43 plants have no developmental defects (30). This result suggested that both the proteins might have some degree of functional overlap, which could be explained by the similar domain and structural organization of the two proteins. Recently it has been reported that the two nucleoplasmin paralogues in *D. melanogaster*, namely DmNLP and DmNPH (nucleophosmin), can form hetero-oligomers, thereby possibly providing additional functional advantages (18). Therefore, similar to these *D. melanogaster* nucleoplasmins, both AtFKBP43 and AtFKBP53 may form a hetero-oligomer to gain additional functional advantages. Such a possibility needs to be verified further.

## MATERIALS AND METHODS

### Generation of bacterial expression constructs

The codon-optimized gene for full-length AtFKBP43 was obtained from Genscript (Piscataway, NJ). Further, bacterial expression constructs of AtFKBP43 N-terminal domain (NTD), spanning residues 1-96, 1-110, 1-136, and 1-164, were individually cloned in a pET22b(+) vector to be expressed with a non-cleavable C-terminal hexahistidine tag. Similarly, an expression construct for AtFKBP43 C-terminal domain (CTD) spanning residues 383-499 was cloned in a pGEX-6P-1 to be expressed with a cleavable N-terminal GST tag. All the clones were confirmed using DNA sequencing.

### Protein overexpression and chromatographic purification

AtFKBP43 NTD and CTD protein expression and purification were performed as described previously (29). Briefly, the AtFKBP43 NTD constructs in a pET22b(+) vector were transformed into *E. coli* BL21 (DE3) cells and allowed to grow at 37 ºC until the OD_600_ value reached 0.5. The cells were induced with 0.2 mM isopropyl β-D-thiogalactopyranoside (IPTG) and incubated at 18 ºC on the shaker for 16 hrs. The cells were subsequently harvested by centrifugation and resuspended in a lysis buffer followed by lysis using an ultrasonic processor from Sonics (Newtown, CT) and centrifuged at a speed of 39,000 xg for 60 mins at 4 °C. The supernatant thus obtained was passed through HisTrap FF 5 ml nickel affinity column (Cytiva, Marlborough, MA). The His-tagged protein bound to the column was eluted with elution buffer using a linear gradient. Size exclusion chromatography was subsequently performed at 4 ºC on the individually eluted proteins for further purification. A HiLoad 16/600 Superdex 200 prep grade column (Cytiva) was used for the purpose in a buffer containing 20 mM Tris (pH 7.5), 150 mM NaCl, 1 mM PMSF, 1 mM EDTA, and 1 mM DTT.

### Analytical size-exclusion chromatography

A Superdex 200 10/300 GL column (Cytiva) was used to perform the analytical SEC experiments at 4 ºC. To analyze the oligomeric status of AtFKBP43 NTDs, the purified protein samples at 0.5 mg/ml concentration were subjected to SEC buffer comprising 20 mM Tris (pH 7.5), 1 mM β-mercaptoethanol, and 300 mM NaCl.

### Analytical ultracentrifugation

An Optima AUC analytical ultracentrifuge (Beckman Coulter, Brea, CA) was used to carry out all the sedimentation velocity analytical ultracentrifugation (SV-AUC) experiments to analyze the molecular mass of AtFKBP43 NTDs and their histone oligomer complexes. AUC experiments for AtFKBP43 (1-96), AtFKBP43 (1-136) and AtFKBP43 (1-164) and their histone oligomer complexes having an OD_280_ ranging from 0.2 to 0.6, were performed as has been described previously (51). The density and viscosity of the sample buffer and the partial specific volume of the protein samples were estimated using SEDNTERP (52). Data fitting was done with a continuous size distribution model, using SEDFIT (53), and the figures were prepared using GUSSI (52).

### Small-Angle X-ray Scattering (SAXS)

SAXS data for AtFKBP43 NTDs and AtFBKBP43 (1-136) histone oligomer complexes were collected at the BM29 BioSAXS beamline of the European Synchrotron Radiation Facility (ESRF, Grenoble, France) on a Pilatus 1M detector. The sample preparation and data collection parameters are given in **Table 1**. SAXS data averaging and buffer subtraction were performed using PRIMUS (36). Data processing was carried out using the ATSAS program (36) and the parameters such as radius of gyration (Rg), maximum particle dimension (Dmax), pair distribution function (P(r)), and the excluded particle volume (Vp) were obtained. The low-resolution envelope structures in P1 and/or P5 symmetries were generated using DAMMIF (54) and averaged using DAMAVER (55). GalaxyHomomer online server (56) was used to generate the structural model for pentameric AtFKBP43 (1-136) and (1-164) with the C-terminal acidic stretches. The low-resolution envelope model of AtFKBP43 NTDs with experimental scattering curves was compared with the calculated scattering curves from the crystal structure using FoXS online server (57). The structure models for AtFKBP43-histone complexes were obtained from the FoXSDock online server (57) using the crystal structures of H2A/H2B and H3/H4 (PDB id: 1KX5). The images of the low-resolution envelope and its overlap with crystal structure were rendered using the PyMOL (Schrödinger, LLC).

### Isothermal titration calorimetry

A MicroCal PEAQ-ITC machine (Malvern Panalytical, Malvern, UK) was used to carry out all isothermal titration calorimetry (ITC) experiments at 25 ºC. AtFKBP43 NTDs, H2A/H2B and H3/H4 were dialyzed at 4 °C with three changes of a buffer comprising 20 mM PIPES (pH 7.4), 1 mM β-mercaptoethanol, and 300 mM NaCl, before the titration experiments. Histone oligomers were taken in the reaction cell at a concentration of 10-15 µM and titrated with 150 µM of individual AtFKBP43 NTDs taken in the injection syringe. Control experiments without the histone complexes were performed for subtraction. The thermodynamic parameters were determined by fitting the isotherms using a one-site binding model with the help of the PEAQ-ITC analysis software. Each titration experiment was repeated thrice for concurrence and data from one experiment has been represented.

### Crystallization, data collection, and data processing

AtFKBP43 (1-110) at a concentration of 22 mg/ml in a buffer containing 20 mM Tris (pH 7.5), 150 mM NaCl, 1 mM EDTA, 1 mM DTT, and 1 mM PMSF was used for crystallization screening. A single rectangular shaped crystal obtained from an optimized condition having 0.1 M Tris (pH 8.5), 25% (w/v) polyethylene glycol 3,350, and 0.2 M ammonium acetate was used for data collection. The crystal was flash-cooled in liquid nitrogen with 20% polyethylene glycol 400 added to the crystallization condition. A high-resolution dataset was collected at the beamline ID23-1 of ESRF (Grenoble, France) and recorded on an Eiger X 4M detector from Dectris. The diffraction data were processed using XDS (58) followed by AIMLESS (59) from the CCP4 suite (60).

### Structure determination

The crystal structure of AtFKBP43 NTD was solved by MolRep program (61) from the CCP4 suite, using the crystal structure of a monomer of AtFKBP53 NTD as a search model (PDB ID: 6J2Z) (29). Crystallographic refinement and model building were conducted using COOT (62) and Phenix refine (63). The stereochemical quality of the structure was checked using the MolProbity program (64), and the structure coordinates have been deposited to PDB with the accession code ***7WIM***. Out of the four pentamers, one had better electron density for the loop regions and was used for preparing figures and structure comparisons. PyMOL was used to prepare the structure figures and structural superpositions.

## ACKNOWLEDGEMENTS

The authors would like to thank their colleagues Dr. Ashish Kumar and Dr. Chinmayee Mohapatra for their help with XRD and SAXS data collections at ESRF and Dr. Deepak T. Nair (RCB, Faridabad) for arranging and coordinating the ESRF trips. The help of their colleague Dr. Aritreyee Datta in editing and finalizing this manuscript is also greatly appreciated.

## DATA AVAILABILITY STATEMENT

The coordinates and structure factors for AtFKBP43 NTD have been deposited at the Protein Data Bank under the entry code ***7WIM***. Scattering data and structural envelopes for AtFKBP43 (1-94), AtFKBP43 (1-136), AtFKBP43 (1-164), AtFKBP43 (1-136)-H2A/H2B and AtFKBP43 (1-136)-H3/H4 have been deposited at the Small Angle X-ray Scattering Biological Data Base (SASBDB) under the accession code ***SASDNN3, SASDNQ7, SASDNT7, SASDNR7* and *SASDNS7***, respectively.

## AUTHOR CONTRIBUTIONS

A.K.S. and K.S. carried out all the experiments, analyzed the data, and wrote the manuscript. S.B. performed part of the histone work, AUC, and SEC experiments. D.V. conceived the project, procured funding, planned and guided the experiments, analyzed the data, and wrote the manuscript.

