## Supplementary Data for "The plant nucleoplasmin AtFKBP43 needs its extended arms for histone interaction"

### SUPPLEMENTARY FIGURES

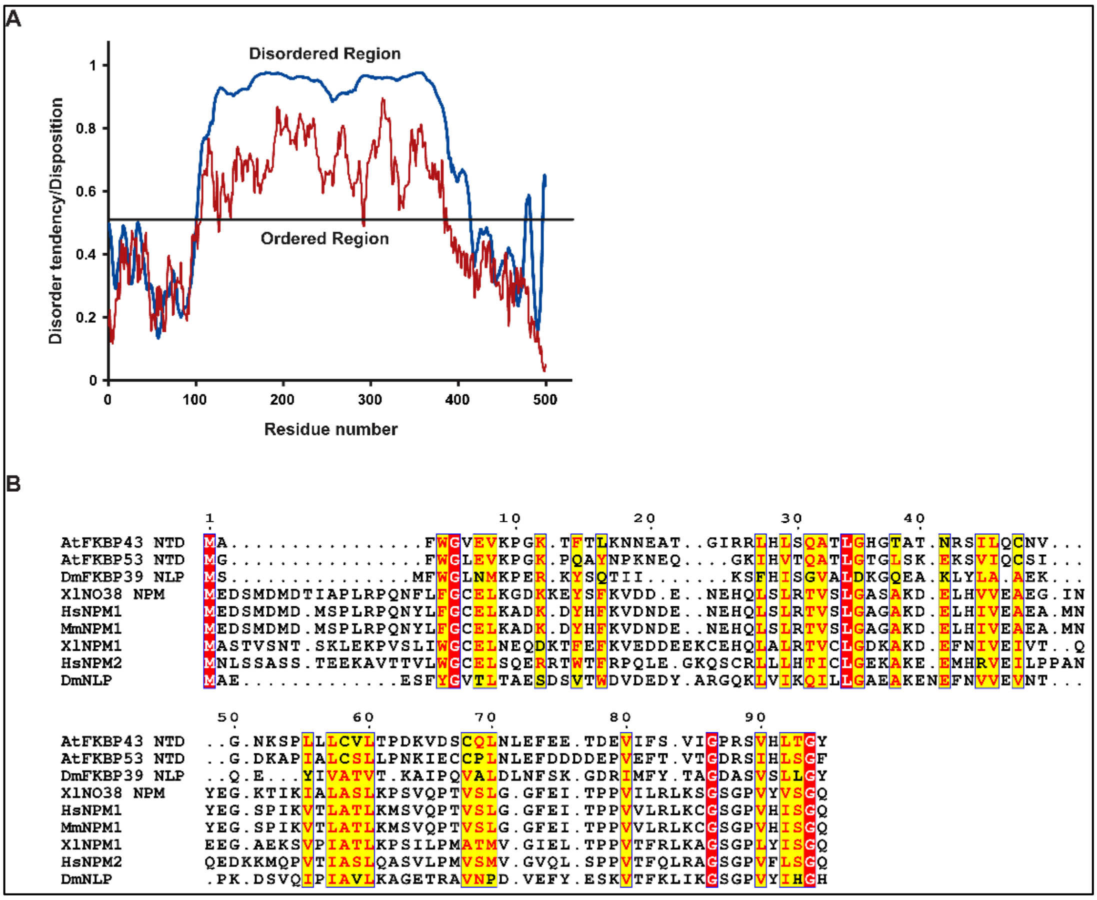


**Figure S1. AFKBP43 disordered region prediction and sequence comparison of AtFKBP43 NTD with other nucleoplasmins. (A)** A graphical representation of the ordered and disordered regions in AtFKBP43, measured by analyzing its sequence using IUpred (dark red) and PONDR-VSL2 (blue). **(B)** Multiple sequence alignment was done using the T-Coffee alignment program for AtFKBP43 NTD with other related proteins such as AtFKBP53 NTD, DmFKBP39 NLP, XlNPM, HsNPM1, MmNPM1, HsNPM2, XlNO38 NPM, and DmNLP. The residue numbering is based on AtFKBP43 NTD. The red-coloured boxes highlight strictly conserved residues, and the yellow-coloured boxes highlight semi-conserved residues.

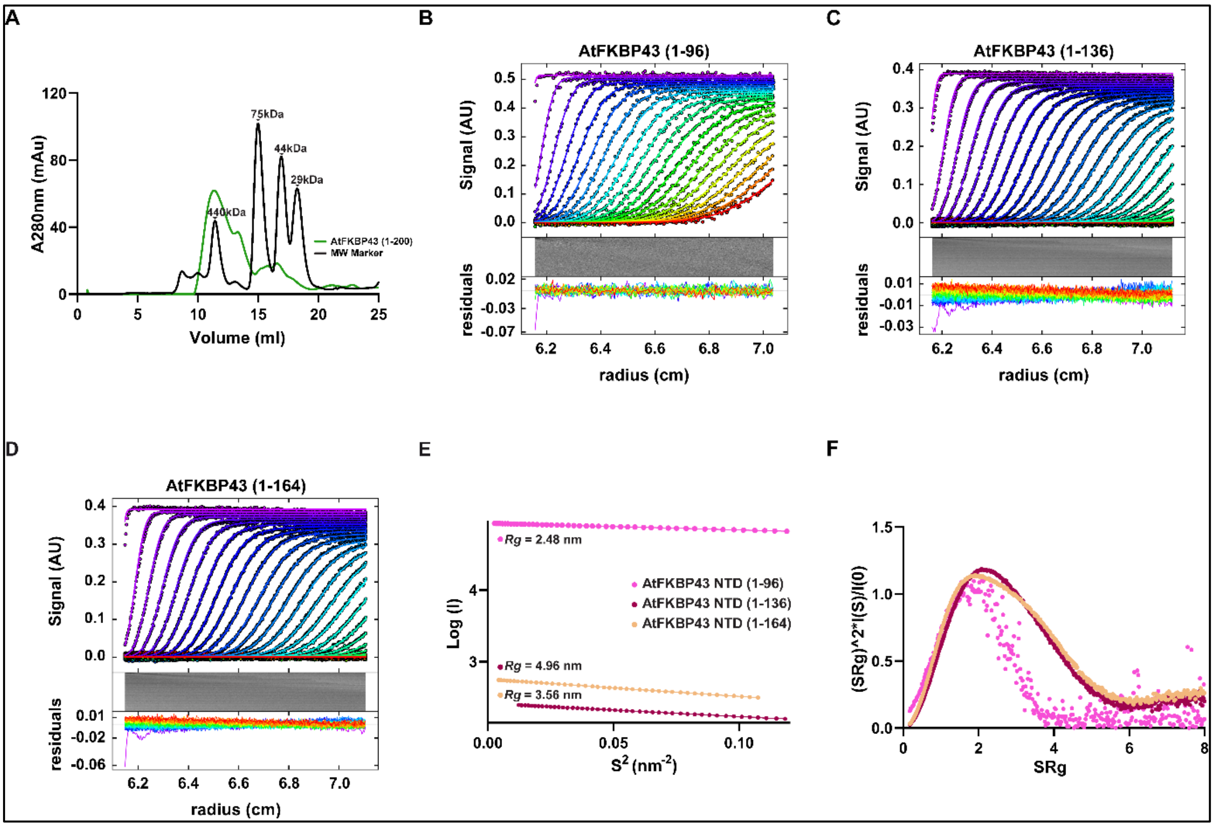


**Figure S2. Analytical-SEC, SV-AUC, and SAXS analyses of AFKBP43 NTDs. (A)** Analytical SEC profiles of AtFKBP43 (1-200). The black curve represents the standard globular proteins of known molecular weight (400kDa - Ferritin, 75 kDa - Carbonic anhydrase, 44kDa - Ovalbumin, 29 kDa - Conalbumin). Superdex 200 10/300 GL column was used for the analytical-SEC experiment. The SV-AUC experiment shows an overlay of the fitted to the experimental curve (upper panel) and the residual plot (lower panel) for **(B)** AtFKBP43 (1-96), **(C)** AtFKBP43 (1-136), and **(D)** AtFKBP43 (1-164). SAXS profile showing **(E)** Guinier plot and **(F)** Kratky plot for AtFKBP43 (1-96) in pink, AtFKBP43 (1-136) in light brown, and AtFKBP43 (1-164) in dark brown colour.

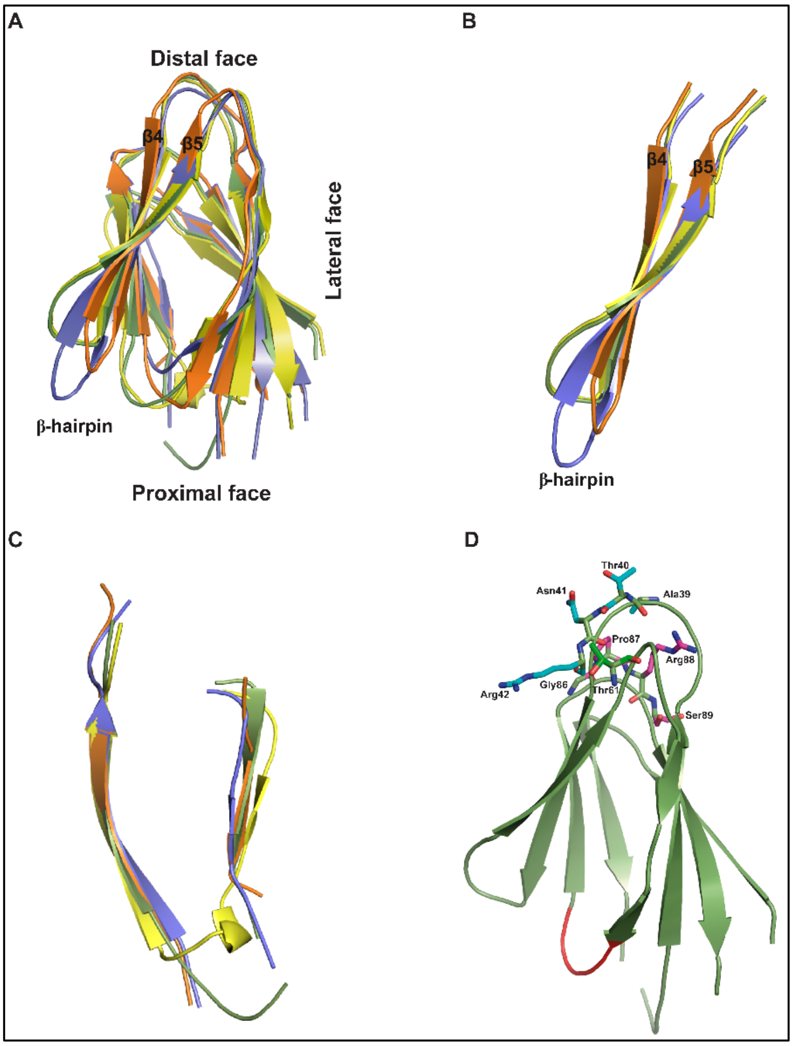


**Figure S3. Structural comparison of AtFKBP43 NTD with other nucleoplasmins. (A)** Structural alignment of AtFKBP43 NTD monomer with AtFKBP53 NTD, DmNLP, and HsNPM1. **(B)** Structural alignment of the β-hairpin motif of AtFKBP43 NTD (green) with that of AtFKBP53 NTD, DmNLP, and HsNPM1. **(C)** Structural alignment of the β2-β3 strand region of AtFKBP43 NTD with that of AtFKBP53 NTD, DmNLP, and HsNPM1. **(D)** AtFKBP43 NTD monomer structure, highlighting in stick model the residues Ala39, Thr40, Asn41 and Arg42 (ATNR motif), Gly86, Pro87, Arg88 and Ser89 (GPRS motif) and Thr61, which would not favour decamerization. AtFKBP43 NTD is shown in green, AtFKBP53 NTD in yellow, DmNLP in orange, and HsNPM1 in blue throughout.

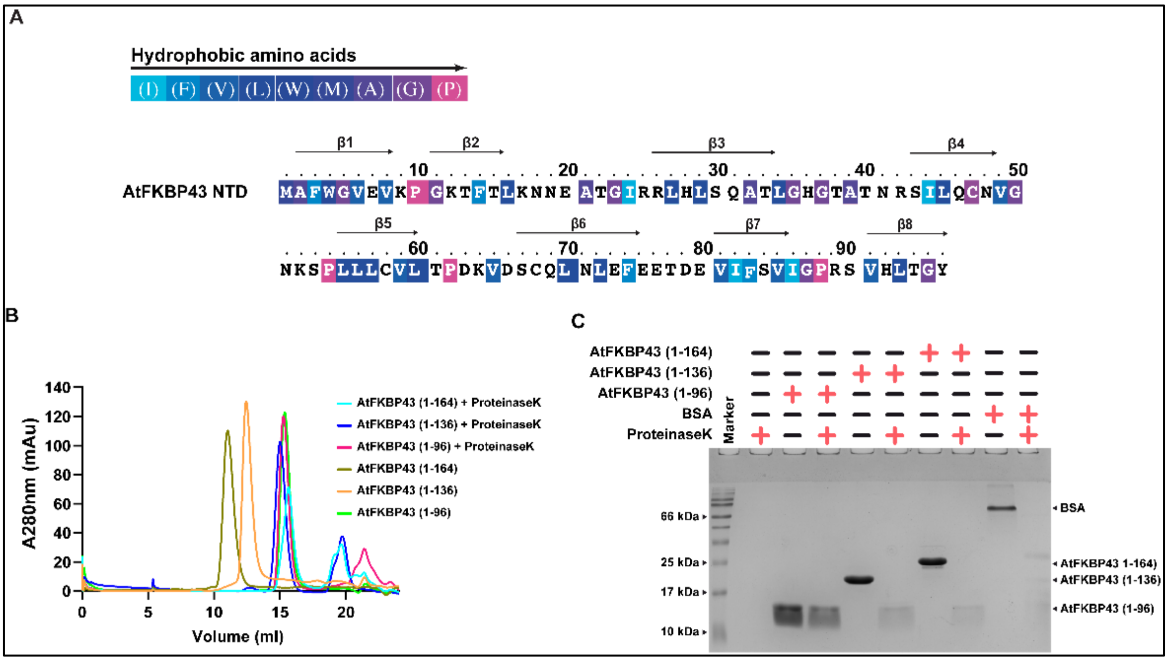


**Supplementary Figure S4. Hydrophobic residue distribution and protease resistance feature of AtFKBP43 NTD. (A)** AtFKBP43 NTD sequence showing the distribution of apolar/hydrophobic residues. The colour code on top explains the degree of hydrophobicity of the residues. **(B)** Analytical size-exclusion chromatogram of AtFKBP43 (1-96), (1-136), and (1-164) in their pentameric form with and without proteinase K treatment. **(C)** 18% SDS-PAGE of the eluted fractions from the analytical SEC. BSA was used as a positive control for proteinase K activity.

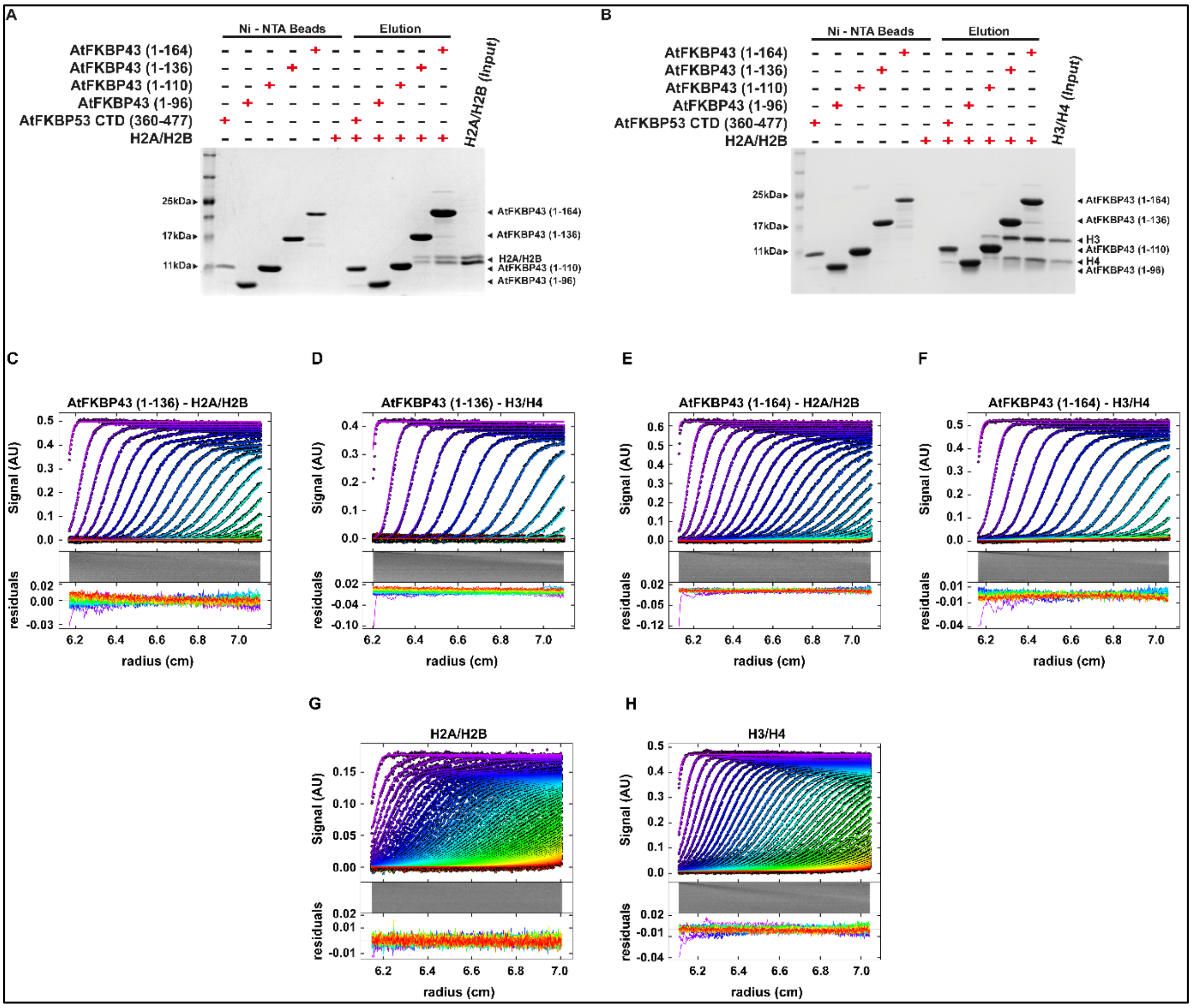


**Supplementary Figure S5. Pull-down assay and SV-AUC profile of AtFKBP43 NTDs with histone H2A/H2B and H3/H4. (A)** The fractions from pull-down assay for AtFKBP43 NTDs with H2A/H2B analyzed on an 18% SDS PAGE gel showing that AtFKBP43 (1-136) and AtFKBP43 (1-164) can bind to H2A/H2B, while AtFKBP43 (1-96) and AtFKBP43 (1-110) do not show any interaction with H2A/H2B **(B)** The fractions from the pull-down assay for AtFKBP43 NTDs with H3/H4 run on an 18% SDS PAGE gel showing that AtFKBP43 (1-136) and AtFKBP43 (1-164), both can bind efficiently to H3/H4, whereas, AtFKBP43 (1-110) can bind less efficiently with H3/H4 and AtFKBP43 (1-96) does not interact with H3/H4. The CTD of AtFKBP53 (residues 360-477) was used as a negative control. The SV-AUC experiment showing an overlay of the fitted and experimental curve (upper panel), and the residual plot (lower panel) for **(C)** AtFKBP43 (1-136)-H2A/H2B, **(D)** AtFKBP43 (1-136)-H3/H4, **(E)** AtFKBP43 (1-164)-H2A/H2B, **(F)** AtFKBP43 (1-164)-H3/H4, **(G)** H2A/H2B, **(H)** H3/H4.

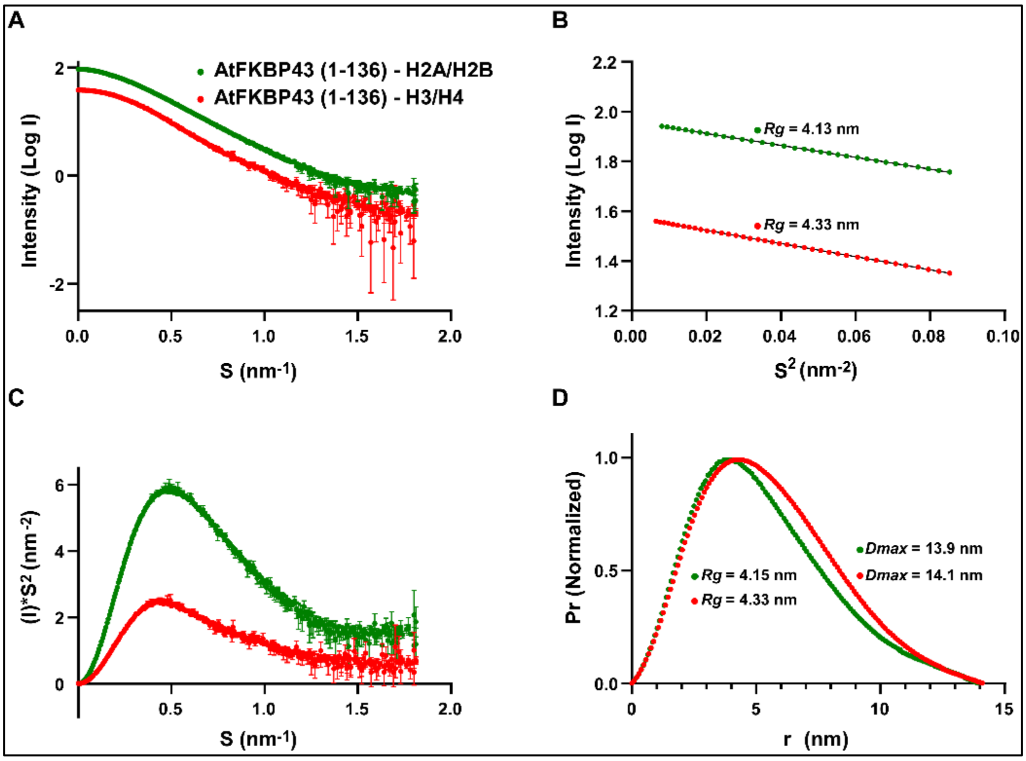


**Figure S6. Small-angle X-ray scattering of AtFKBP43 (1-136)-H2A/H2B and AtFKBP43 (1-136)-H3/H4 complexes.** All the profiles for AtFKBP43 (1-136)-H2A/H2B have been presented in green and those for AtFKBP43 (1-136)-H3/H4 in red. **(A)** Logarithmic plot showing the relative scattering intensity profile of AtFKBP43 (1-136) histone complexes as a function of scattering vector s $s = 4\pi\sin(\theta)/\lambda$, where $2\theta$ is the scattering angle, and $\lambda$ is the wavelength. **(B)** Guinier plot showing the linear fit of the experimental data in the low Q region. **(C)** Kratky plot for AtFKBP43 (1-136) histone complexes. **(D)** Paired distance distribution functions plot for AtFKBP43 (1-136) histone complexes showing logarithmic scattering intensity as a function of momentum transfer.

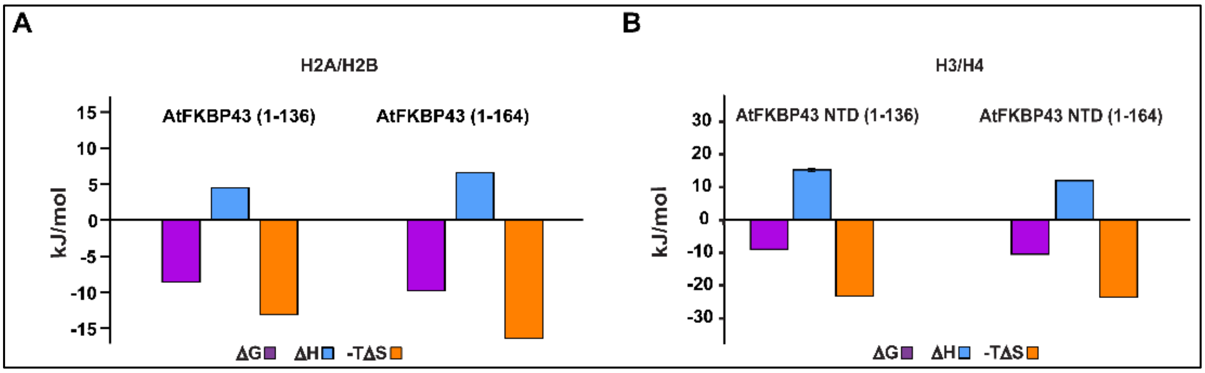


**Figure S7. ITC thermodynamic parameters of AtFKBP43 (1-136) and AtFKBP43 (1-164) interaction with histone oligomers. (A)** Bar graph representing ΔG, ΔH, and TΔS for AtFKBP43 (1-136) and AtFKBP43 (1-164) interaction with H2A/H2B. **(B)** Bar graph representing ΔG, ΔH, and TΔS for AtFKBP43 (1-136) and AtFKBP43 (1-164) interaction with H3/H4.

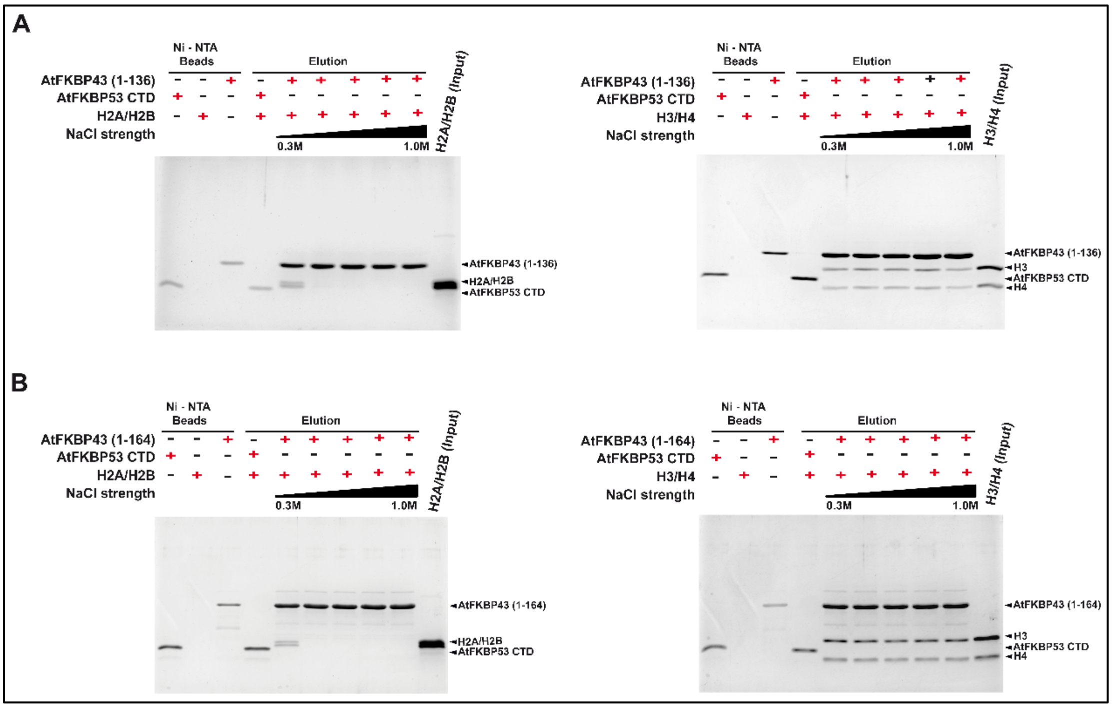


**Figure S8. Stability assay for AtFKBP43 (1-136) and AtFKBP43 (1-164) complexes with H2A/H2B and H3/H4. (A)** Eluted fractions from separate pull-down experiments of AtFKBP43 (1-136)-H2A/H2B complex (left panel) and AtFKBP43 (1-136)-H3/H4 complex (right panel), performed with increasing salt concentrations (0.3 M to 1.0 M) analyzed on an 18% SDS-PAGE gel. **(B)** Eluted fractions from separate pull-down experiments of AtFKBP43 (1-164)-H2A/H2B complex (left panel) and AtFKBP43 (1-164)-H3/H4 complex (right panel), performed with increasing salt concentrations (0.3 M to 1.0 M) analyzed on an 18% SDS-PAGE gel. The presence of H3/H4 bound to both the lengths of AtFKBP43, even at higher ionic strength conditions, suggests a major role for hydrophobic interactions in stabilizing both the complexes; whereas, it appears to be more electrostatic for H2A/H2B.

#### SUPPLEMENTARY TABLES

**Table S1: Secondary structure elements estimated from AtFKBP43 NTDs samples using Circular Dichroism**

| | Helix | Turn | Strand | Unstructured | NRMSD |
|---|---|---|---|---|---|
| AtFKBP43 (1-96) | 3% | 43% | 19% | 35% | 0.03 |
| AtFKBP43 (1-136) | 8% | 21% | 21% | 50% | 0.02 |
| AtFKBP43 (1-164) | 9% | 20% | 18% | 53% | 0.03 |

**Table S2: Melting temperature (*T*m) of AtFKBP43 (1-96) samples as estimated from thermal Circular Dichroism experiments**

| **Analyte** | ***T*m (°C)** |
|---|---|
| AtFKBP43 (1-96) + 4 M Urea | 37.09 ± 0.78 |
| AtFKBP43 (1-96) + 3 M Urea | 46.95 ± 0.45 |
| AtFKBP43 (1-96) + 2 M Urea | 58.02 ± 0.18 |
| AtFKBP43 (1-96) + 1 M Urea | 62.31 ± 0.17 |
| AtFKBP43 (1-96) control | 63.47 ± 0.24 |
| AtFKBP43 (1-96) + 0.25 M NaCl | 64.37 ± 0.20 |
| AtFKBP43 (1-96) + 0.5 M NaCl | 68.03 ± 0.22 |
| AtFKBP43 (1-96) + 1 M NaCl | 72.47 ± 0.28 |
| AtFKBP43 (1-96) + 2 M NaCl | 84.16 ± 2.25 |
| AtFKBP43 (1-96) + 3 M NaCl | 93.28 ± 4.96 |
| AtFKBP43 (1-96) + 4 M NaCl | 89.51 ± 1.59 |
| AtFKBP43 (1-96) + 5 M NaCl | 107.35 ± 3.27 |

#### SUPPLEMENTARY METHODS

##### *In silico* analysis

The web servers IUpred (1, 2) and PONDR-VSL2 (3) with default settings were used to analyze the amino acid sequence of AtFKBP43 to identify the ordered and disordered regions.

##### Circular dichroism spectroscopy

Circular dichroism (CD) spectra were recorded in the far-ultraviolet (UV) region of 190-260 nm using a J-1500 CD spectrophotometer (Jasco, Tokyo, Japan). Spectra were acquired for purified AtFKBP43 (1-96) at a concentration of 0.25 mg/ml in a buffer containing 50 mM sodium phosphate (pH 7.5) and 1 mM DTT, taken in a quartz cuvette of 1.0 mm path length. The spectra were analyzed using the online program BeStSel (4) to obtain the percentage of secondary structural elements present in the protein. CD thermal denaturation experiments were also carried out, wherein the CD signal of the protein was monitored at a wavelength of 222 nm from 25 °C to 95 °C at a thermal ramping speed of 4 °C/min. Similarly, the CD signal at 222 nm was collected for the protein incubated for 30 minutes at room temperature with urea in a range of concentrations varying from 0 to 4 M and with NaCl in a range of concentrations varying from 0.25 M to 5 M, separately and the thermal denaturation of the treated protein samples monitored. The thermal denaturation data was collected by examining the CD spectra as a function of temperature. The CD spectra values were normalized (from 0 to 1) to simplify the overlapped curves using the software CDpal (5). The data were fitted to a two-state model (fully irreversible N to D transition), and the melting temperature (Tm) was determined for all the samples.

##### Protease stability assay

For protease stability assay, 1 mg of the different AtFKBP43 NTD stretches after purification were incubated separately with a 50:1 ratio of proteinase K in a buffer containing 20 mM Tris (pH 7.5), 1 mM β-mercaptoethanol, and 300 mM NaCl, followed by incubation at 4 ºC for 30 mins. Subsequently, analytical SEC was performed on the individual proteinase K-treated samples using a Superdex 200 10/300 GL column at 4 ºC, and the eluted fractions were run on an 18% SDS-PAGE gel and analyzed after Coomassie Blue staining.

##### Expression and purification of core histones

The codon-optimized constructs for the core histones H2A, H2B, H3, and H4 were a kind gift from Dr. Curtis A. Davey (Nanyang Technological University, Singapore). These core histones were expressed without a fusion tag as described before (6). Briefly, the constructs in the pET21a (+) vector were transformed individually into *E. coli* BL21 (DE3) pLysS cells and subsequently grown in 2xYT medium at 37 ºC. The histones were overexpressed by induction with 0.4 mM IPTG when the $OD_{600}$ of the culture reached 0.4, and the induction was allowed to proceed for 3 hrs at 37 ºC. A HiLoad 16/600 Superdex 200 prep grade column was used to perform SEC under denaturing conditions to purify the individual histones obtained in the inclusion bodies. The histones thus purified were subjected to dialysis against MilliQ water (Merck-Millipore, Burlington, MA), aliquoted into smaller volumes, lyophilized, and stored in a -80 ºC freezer.

**Preparation of histone oligomers**

Histone oligomers were prepared as described previously.[6] The lyophilized core histones H2A, H2B, H3 and H4 dissolved in a denaturing buffer containing 7 M Guanidinium hydrochloride were mixed in partners (H2A and H2B or H3 and H4) in equimolar stoichiometry and refolded to obtain H2A/H2B dimer and H3/H4 tetramer by dialyzing against a refolding buffer comprised of 20 mM Tris (pH 7.5), 10 mM β-mercaptoethanol, 2 M NaCl, 1 mM PMSF, and 1 mM EDTA. The obtained histone oligomers were purified by SEC using a HiLoad 16/600 Superdex 200 prep grade column, pre-equilibrated with the refolding buffer. The peak fractions were pooled together, concentrated, and used for the various experiments.

**Pull-down assay**

The various stretches of purified AtFKBP43 NTD with His-tag at a concentration of 5 μM were individually mixed with reconstituted H2A/H2B or H3/H4 histone oligomers (20 μM) in 300 μl of a binding buffer containing 20 mM Tris (pH 7.5), 30 mM imidazole, 300 mM NaCl, 1 mM β-mercaptoethanol, and 10 μg/ml BSA. The individual mixtures were incubated at 4 ºC for 30 min with Ni-NTA agarose beads (Invitrogen, Waltham, MA), pre-equilibrated with the binding buffer. After incubation, the beads were washed extensively with a wash buffer containing 20 mM Tris (pH 7.5), 50 mM imidazole, 300 mM NaCl, 1 mM β-mercaptoethanol, and 0.2% Tween 20. Next, the bound protein complex was eluted in a buffer comprising 20 mM Tris (pH 7.5), 500 mM imidazole, 300 mM NaCl, and 1 mM β-mercaptoethanol. Elution fractions were run on an 18% SDS-PAGE gel and stained with Coomassie Blue for analysis.

The stability of the complexes of AtFKBP43 NTDs with H2A/H2B and H3/H4 in the presence of salt was also studied using pull-down assay. Towards this end, the complexes were incubated in individual reactions with buffers having increasing concentrations of NaCl, in the range of 0.3 M to 1 M. Further, the beads were washed, and the proteins eluted and run on an18% SDS-PAGE gel for analysis.
